# Synthesis of low immunogenicity RNA with high-temperature *in vitro* transcription

**DOI:** 10.1101/815092

**Authors:** Monica Z. Wu, Haruichi Asahara, George Tzertzinis, Bijoyita Roy

**Affiliations:** New England Biolabs, Inc, Ipswich, Massachusetts, 01938, United States of America

## Abstract

The use of synthetic RNA for therapeutics requires that the *in vitro* synthesis process be robust and efficient. The technology used for the synthesis of these *in vitro-*transcribed mRNAs, predominantly using phage RNA polymerases (RNAPs), is well established. However, transcripts synthesized with RNAPs are known to display an immune-stimulatory activity *in vivo,* that is often undesirable. Previous studies have identified double-stranded RNA (dsRNA), a major by-product of the *in vitro* transcription (IVT) process, as a trigger of cellular immune responses. Here we describe the characterization of a high-temperature IVT process using thermostable T7 RNAPs to synthesize functional mRNAs that demonstrate reduced immunogenicity without the need for a post-synthesis purification step. We identify features that drive the production of two kinds of dsRNA by-products—one arising from 3’ extension of the run-off product and one formed by the production of antisense RNAs—and demonstrate that at a high temperature, T7 RNAP has reduced 3’-self extension of the run-off product. We show that template-encoded poly-A tailing does not affect 3’-self extension but reduces the formation of the antisense RNA by-products and that combining high-temperature IVT with template-encoded poly-A tailing prevents formation of both kinds of by-products.

## INTRODUCTION

One prominent use of *in vitro* transcription in the past few years has been to generate mRNAs for biopharmaceutical and therapeutic applications. The technology used for the synthesis of these *in vitro* transcribed mRNAs—predominantly using phage RNA polymerases (RNAPs)—is robust and well established for the large-scale production of synthetic RNA. However, it is also known that introduction of synthetic *in vitro* transcribed mRNAs into cells or animal models results in an immune response against the synthetic molecules [1–5]. Such outcomes are undesirable in therapeutic applications in which an immune response is detrimental or unnecessary (for example, protein-replacement therapies). Incorporation of modified nucleosides into synthetic mRNA mitigates the immune response to some extent by mimicking endogenous mRNAs [4, 6–9]. Another major stimulant of the immune response comes from contaminants present in the *in vitro* transcription reactions. A major byproduct identified in *in vitro* transcription (IVT) reactions is dsRNA; this can arise from T7 RNAP’s RNA-dependent RNA polymerase activity [10–16]. Introduction of transcripts synthesized using T7 RNAP has been shown to activate cytosolic sensors, such as RIG-I and MDA5, that activate the innate immune system in response to viral double-stranded RNAs (dsRNAs) [17–19]. Recent studies have identified two main types of byproducts in the IVT reaction that result in formation of dsRNA molecules. The first is formed by 3’-extension of the run-off products annealing to complementary sequences in the body of the run-off transcript either in *cis* (by folding back on the same RNA molecule) or *trans* (annealing to a second RNA molecule) to form extended duplexes [11, 20]. The second type of dsRNA molecules is formed by hybridization of an antisense RNA molecule to the run-off transcript. The antisense RNA molecules have been reported to be formed in a promoter- and run-off transcript-independent manner [21]. The mechanistic and structural requirements for formation of these byproducts are not well understood, and therefore, it is a challenge to devise methods to prevent their formation or enable their removal.

Because the workflow for therapeutic RNA synthesis has to be compatible with large scale up processes, preventing the formation of these dsRNA byproducts in the reaction is more desirable than adding a post-synthesis purification step, such as chromatography-based purification approaches [22–24]. Here we describe a simple and scalable method for preventing the synthesis of dsRNA byproducts in the IVT reaction. We describe the use of thermostable T7 RNAPs and the effect of the temperature of the *in vitro* transcription reaction in the formation of dsRNA byproducts. We characterize the dsRNA products in the IVT reactions and show that high-temperature transcription results in reduction of 3’-extended RNA, but not antisense RNA byproducts. We also demonstrate that the presence of a template-encoded poly-A tail has a beneficial effect on the formation of antisense byproducts but not on 3’-extended byproducts. Finally, we show that mRNAs synthesized with thermostable RNAPs at higher temperatures are functional and have reduced immunogenicity *in vivo*.

## MATERIALS AND METHODS

### Oligonucleotides

All DNA and RNA oligonucleotides for *in vitro* transcription were synthesized by Integrated DNA Technologies (IDT, Coralville IA) and the sequences are available in Supplementary Table S1.

### *In vitro* transcription

For IVT of the shorter oligonucleotides, the forward and reverse oligonucleotides were annealed in DNA annealing buffer (10 mM Tris pH 7.5, 50 mM NaCl, 1 mM EDTA), heated at 95°C for 5 minutes and then cooled to room temperature. Transcription reactions were performed with 5 mM of each ribonucleotide triphosphate. Reactions with wild-type T7 RNAP were carried out in 40 mM Tris-HCl pH 7.9, 19 mM MgCl_2_, 5 mM DTT, 1 mM spermidine and supplemented with RNase inhibitor (1000 units/mL) and inorganic pyrophosphatase (4.15 units/mL) (New England Biolabs) at 37°C for 1 hour. Reactions with thermostable T7 RNA polymerase (TsT7-1 – Hi-T7_TM_ RNA polymerase; New England Biolabs and TsT7-2 – Thermostable T7 RNAP; Toyobo Life Sciences) were carried out in a buffer containing 40 mM Tris-HCl pH 7.5, 19 mM MgCl_2_, 50mM NaCl, 5 mM DTT, 1 mM spermidine at varying temperatures (37°C to 60°C) for 1 hour. Reactions were supplemented with RNase inhibitor (1000 units/mL) and inorganic pyrophosphatase (4.15 units/mL) (New England Biolabs). The *in vitro* transcribed RNAs were treated with Turbo_TM_ DNase (Invitrogen) and purified with the Oligo Clean-Up and Concentration kit (Norgen Biotek Inc). For the 512B transcripts, reactions were cleaned up with the Monarch® RNA Cleanup Kit (New England Biolabs).

IVT of long mRNAs (Cypridina luciferase (CLuc; 1700 nucleotides), red fluorescent protein (RFP; 795 nucleotides) or green fluorescent protein (GFP; 761 nucleotides)) were performed from a linearized plasmid (digested with NotI) or a PCR-amplified linear template. For each reaction, 30 nM DNA template and 30 nM T7 RNAP variants were used. All four nucleoside triphosphates in the reaction, natural or modified, were used at a final concentration of 4 mM each. For generation of nucleoside-modified mRNAs, UTP was replaced with triphosphate-derivative of pseudouridine (Trilink Biotechnologies). A 120-nucleotide poly-A tail was template-encoded unless otherwise indicated. The *in vitro* transcribed RNAs were treated with Turbo_TM_ DNase (Invitrogen) and followed by spin column clean-up (MEGAclear™ Transcription Clean-Up Kit). The mRNAs were post-transcriptionally capped with Vaccinia capping enzyme and treated with mRNA Cap 2’-O-Methyltransferase (New England Biolabs) as recommended by the manufacturer.

### T7 RNA polymerase

Wild type T7 RNAP was purified by Ni-NTA chromatography in 50 mM Hepes KOH pH 7.5, 100 mM NaCl, 10mM DTT, 0.1% Triton X-100 and stored in 50% glycerol. The thermostable T7 RNAPs (TsT7-1 – Hi-T7_TM_ RNA polymerase; New England Biolabs and TsT7-2 – Thermostable T7 RNA polymerase; Toyobo Life Sciences [25–28]) were either purchased from the respective manufacturers.

### Molecular beacon assay for *in vitro* transcription efficiency

Efficiency of *in vitro* transcription reactions was monitored using a modified molecular beacon assay based on [29] performed in 40 mM Tris-HCl pH 7.9, 19 mM MgCl_2_, 50mM NaCl, 5 mM DTT, 1 mM spermidine, 30 nM DNA template, 30 nM T7 RNAP variants, and 0.5 μM molecular beacon probe. The template was a 22 kb plasmid DNA linearized 6 kb downstream of the T7 promoter, where the beacon target sequence occurs immediately upstream of the linearization site. The target sequence is a 24-nucleotide segment complementary to the loop sequence (underlined) of the DNA oligonucleotide molecular beacon: 5’-CCTGC <u>GATT GAA CAC GTG</u> <u>GGT CAG AGA GG</u> GCAGG-3’. The fluorescent dye 6-FAM was conjugated to the 5’ end and the quencher (BHQ1) to the 3’ end (Integrated DNA Technologies, Coralville, IA). Reactions were run at various temperatures in the CFX96 Touch Real-Time PCR Detection System (Bio-Rad, Hercules, CA) for 1 hour. End-point fluorescence units for each polymerase were graphed against temperature.

### Melting temperature and thermostability of RNAP

Wild type T7 RNAP and thermostable T7 variants were diluted to a starting concentration of 2µg in Tris-HCl, pH 7.5 buffer (40mM Tris-HCl, 50mM NaCl, 19mM MgCl_2_, 1mM DTT, 2mM spermidine). Each enzyme was then serially diluted three times at 1:2 ratio for final protein amounts of 2µg, 1µg, 500ng and 250ng. Fluorescence-based thermal unfolding experiments were performed using the Prometheus NT.48 (NanoTemper Technologies). 5µL of each sample was loaded into the machine. The temperature was increased by 2°C/min from 20°C to 80°C and the fluorescence at emission wavelengths of 330nm and 350nm was measured. Final data is displayed as the first derivative of the ratio of 350nm/330 nm emission. Final T_m_ for each enzyme is the average of the four dilutions.

### Intact mass analysis of IVT products

IVT reactions were treated with Turbo_TM_ DNase (Invitrogen) and cleaned up with an Oligo Clean-Up and Concentration kit (Norgen Biotek Inc). 100pmol of each IVT sample was used for intact oligonucleotide mass spectrometry analysis at Novatia, LLC (Newtown, PA, USA) using on-line desalting, flow injection electrospray ionization on a Thermo Fisher Scientific LTQ-XL ion trap mass spectrometer and analyzed with a ProMass Deconvolution software.

### dsRNA immunoblot

Crude IVT reactions or purified IVT RNA samples were spotted onto positively charged nylon membranes (Nytran_R_ SC, Sigma-Aldrich). The membranes were blocked in 5% (w/v) non-fat dried milk in TBS-T buffer (20 mM Tris, pH 7.4, 150 mM NaCl, 0.1 % (v/v) Tween-20). For the detection of dsRNA, the membranes were incubated with J2 anti-dsRNA antibody (1:5000; Scicons) at 4°C overnight. The blots were probed with IR Dye™ -680 or -800 conjugated secondary antibodies (Cell Signaling Technologies). dsRNA ladder (New England Biolabs) and poly(I:C) (Invivogen) were used as positive controls.

### Gel electrophoresis for antisense RNA and 3’ extended by-product detection

The crude IVT reactions or purified transcripts were analyzed in TBE 20% polyacrylamide gel (native condition) or 6-15% TBE-Urea polyacrylamide gels (for denaturing condition). Typically, purified IVT RNA was denatured in 2X RNA loading dye (New England Biolabs) and heated at 70°C for 2 minutes before analyzing on TBE-Urea polyacrylamide gels. SybrGold staining (ThermoFisher Scientific) was performed for RNA visualization and gels were scanned with an Amersham™ Typhoon™ Biomolecular Imager (GE Healthcare).

### Cells

Human embryonic kidney cells HEK293 were obtained from American Type Culture Collection and were cultured in Dulbecco’s modified Eagle’s medium (DMEM) supplemented with L-glutamine (ThermoFisher Scientific) and 10% fetal calf serum. Human dendritic cells were purchased from Lonza Group AG and were differentiated according to the manufacturer’s recommendations.

### HPLC purification of IVT mRNA

CLuc mRNA was purified by high performance liquid chromatography (HPLC) according to a protocol described previously [22] using a linear gradient of 38%-70% buffer B (0.1 M triethylammonium acetate (TEAA), pH 7.0, 25% (v/v) acetonitrile) in buffer A (0.1 M TEAA, pH 7.0) at a flow rate of 5 mL/min. The RNA from the collected fractions was concentrated and desalted with centrifugal filter units (30 kDa MWCO) (Millipore).

### mRNA transfection and expression analyses

Synthesized, purified mRNAs were transfected into human embryonic kidney cells using the TransIT-mRNA transfection kit as recommended by the manufacturer (Mirus Bio). The expression of the CLuc mRNA was analyzed by measuring the luciferase activity from the media at various time points (15, 30, 60, 90, 120, 180, 240, 300, 360 minutes post-transfection). The luciferase activity was measured with BioLux® Cypridina Luciferase Assay Kit (New England Biolabs) using a Centro LB 960 luminometer (Berthold) in relative light units (RLU).

### Immune response assay

Dendritic cells (human) were treated with PBS (ThermoFisher Scientific), R-848 (Invivogen), poly(I:C) (Invivogen), or TransIT-complexed IVT RNA. 24 hours post-transfection, supernatant was harvested and the level of IFN-α was measured as recommended by the manufacturer (PBL Assay Science).

## RESULTS

### 3’-extended byproducts are formed during IVT

To determine the extent and nature of the dsRNA byproducts from IVT reactions, we tested multiple templates of varying lengths and sequences. We used mRNAs encoding for three different proteins (*Cypridina* luciferase (CLuc), red fluorescent protein (RFP), and green fluorescent protein (GFP)). IVT reactions were performed with wild-type T7 RNAP under standard conditions (5 mM each rNTP; 1 hour at 37°C). For detection of the dsRNA byproducts, we utilized a standard immunoblot assay that involves recognition of continuous double-stranded structures (>40 bp in length) by a monoclonal antibody (J2) specific for dsRNA (Supplementary Figure S1)[30, 31]. Consistent with previous reports, the immunoblot assay showed varying levels of dsRNA byproducts in all of the samples, suggesting that under the IVT conditions tested, detectable amounts of dsRNA contaminants are generated (Figure 1A). Although batch-to-batch variation in the relative amount of dsRNA was detected, overall the trend was consistent. Furthermore, the immunoblot signal diminished when the IVT reactions were treated with RNase III, suggesting that it is generated from bona fide dsRNA contaminants (Figure 1A). We therefore established that these templates could be used to evaluate the effects of varying reaction conditions and methods to reduce the formation of these dsRNA byproducts.

**Figure 1.**
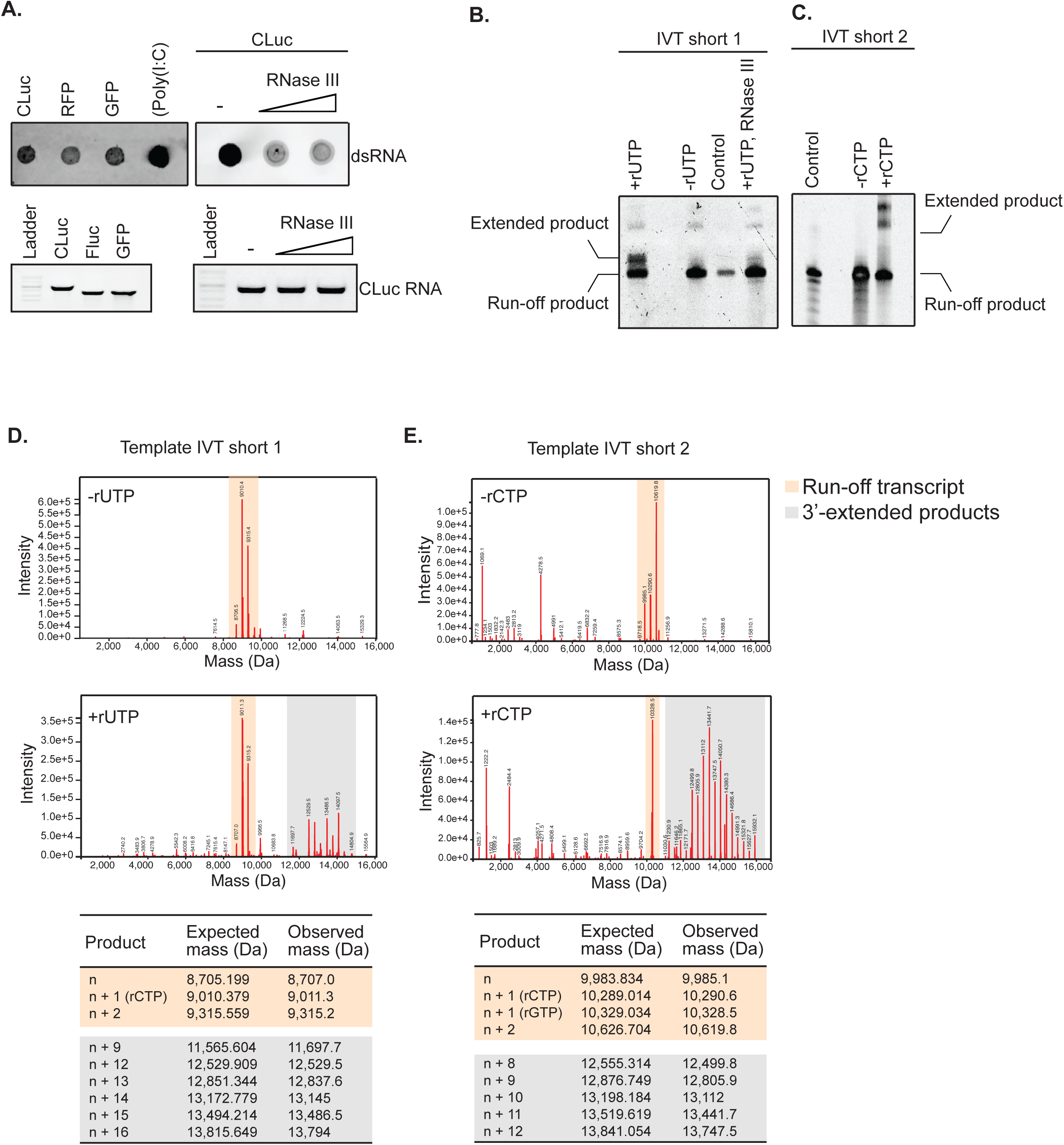
3’ extended RNA byproducts are formed during *in vitro* transcription (IVT) reactions. **A)** dsRNA immunoblot assay using a dsRNA-specific antibody (J2) on three different mRNA sequences (CLuc, RFP and GFP). Poly(I:C) and RNase III treatment is used as a control to detect and validate dsRNA respectively. **B, C)** IVT reactions using short templates (30 bp) ran on denaturing gels under standard conditions (four rNTPs), and under conditions where one rNTP is eliminated to prevent formation of the dsRNA products (-rUTP for **B** and -rCTP for **C**). RNase III digestion control to confirm the dsRNA nature of the extended by-product. Chemically synthesized RNA used as a control. **D, E)** Intact mass spectrometry of IVT products when all four rNTPs are present (+rUTP for **D** and +rCTP for **E**), and when one rNTP is eliminated to prevent formation of the dsRNA products (-rUTP for **D** and -rCTP for **E**). Expected mass of extended products is calculated based on average rNTP values.

Since the immunoblot assay does not distinguish between the structure or origin of these dsRNA byproducts, we established an alternative method for detection of the dsRNA contaminants. We designed a short DNA template (Template IVT short 1; Supplementary Table S1) that was used to synthesize a 30 nucleotide (nt) run-off RNA product and can be subjected to intact mass spectrometry (MS) analysis. Intact MS analysis would allow for the detection of the run-off product, as well as any other spurious products that are present in the IVT reaction and not resolved by standard gel electrophoresis methods. Furthermore, it could also differentiate between the formation of a dsRNA region due to secondary structures of the run-off RNA from an actual by-product that is different from the run-off transcript. The DNA template sequence was designed with only three nucleotides (G, A, and C) so that the IVT reaction could be performed with just three rNTPs and thereby prevent dsRNA formation in the absence of the fourth nucleotide (rUTP) (Supplementary Figure S2A). Addition of the fourth nucleotide (rUTP) allows base-pairing with adenine in the run-off transcript to form dsRNA regions (Supplementary Figure S2A). Analysis of the IVT products using denaturing gel electrophoresis showed the presence of the expected run-off transcript compared to a chemically synthesized RNA used as a control when three NTPs (-rUTP lane) were included in the IVT reaction (Figure 1B). In contrast, two major RNA products were detected when all four NTPs were present in the reaction (Figure 1B; +rUTP lane). The additional extended RNA product detected, migrated at a higher position suggesting that it is greater in length than the expected run-off transcript (Figure 1B). Furthermore, RNase III treatment of the products from the IVT reaction with all four NTPs resulted in the disappearance of the higher molecular weight product, confirming its double-stranded nature (Figure 1B). The IVT reactions were further subjected to intact MS analysis to identify the molecular weight of all RNA products in the reaction. In the –rUTP reactions, we observed the presence of three main RNA species represented by the expected run-off product (8,707 Da), and transcripts that could be attributed to a single nucleotide addition (n+1) to the expected runoff transcript (+C (9,011.3 Da)) and some di-nucleotide additions (n+2; +CG (9,315.2 Da)) (Figure 1D). The synthesis of the non-templated n+1 and n+2 products in IVT reactions has been reported previously and is thought to be an inherent property of T7 RNAP [32, 33]. In contrast, in the transcription reactions with all four NTPs (+rUTP reactions), in addition to the n, n+1, n+2 products, we detected a heterogeneous distribution of RNA products which were up to 22-nt longer (>14,804.9 Da) than the expected 30-nt run-off product (Figure 1D). RNase III treatment followed by intact mass analysis resulted in the loss of the higher-molecular-weight RNA products but not the n, n+1, and n+2 RNA species (Supplementary Figure S3). In order to rule out potential sequence-specific effects, the same set of experiments was repeated with another template (sequence consisting of G, A, T; Template IVT short 2; Supplementary Figure S2B), where rCTP could be omitted from the reaction to form the run-off transcript and dsRNA would be formed only when rCTP was added to the reaction (Figure 1C, E). The same pattern was observed with this second template (Figure 1C, E). Altogether, the intact mass and denaturing gel analyses of the short IVT templates suggest that during IVT spurious, 3’-extended products are synthesized in the reaction and some of these can form dsRNA structures, supported by their RNase III sensitivity. These observations are consistent with a recent study that demonstrated the synthesis of a heterogeneous population of 3’-extended RNA products during IVT from a completely unrelated DNA template [20].

Taken together, these results demonstrate that intact mass analysis of short template IVT products can be used together with immunoblot analysis of longer mRNAs to evaluate and monitor dsRNA contaminants in IVT reactions.

### Thermostable T7 RNAPs can synthesize RNA efficiently

Because wild-type T7 RNAP has the propensity to make spurious 3’-extended byproducts, we investigated whether changing the reaction conditions could alter the formation of these byproducts. One of the proposed mechanisms for formation of 3’-extended byproducts is the rebinding of the run-off transcript by T7 RNA polymerase followed by self-priming [13, 20]. We hypothesized that altering the reaction temperature could prevent rebinding of the RNA polymerase to the run-off transcript and thereby reduce the formation of the 3’-extended contaminants. To test this hypothesis, we focused on commercially available thermostable (Ts) RNAPs (TsT7-1 and TsT7-2; Supplementary Figure S4), which allowed us to raise the temperature of the IVT reaction. As a first step, we evaluated the thermostability of the RNAPs and the optimum temperature for the IVT reactions. As compared to wild-type T7, both TsT7-1 and TsT7-2 had higher thermostability as indicated by the higher melting temperature of the proteins determined using nano-differential scanning fluorimetry (nano-DSF) (Figure 2A). The TsT7 RNAPs were transcriptionally active at temperatures greater than 45°C, where the activity of wild-type T7 RNAP is compromised, as analyzed with a molecular beacon assay that measures the real-time synthesis of the RNA in the reaction (Figure 2B) as well as with gel electrophoresis analysis of the CLuc mRNA product (Figure 2C). Even though TsT7-1 was active at 60°C, high-temperature IVT reactions were performed at 50°C to keep the reaction conditions identical between the two TsT7 polymerases. Importantly, the integrity and yield of the RNAs synthesized from both TsT7 RNAPs were comparable for the templates tested (CLuc; Figure 2C; Supplementary Figure S5).

**Figure 2.**
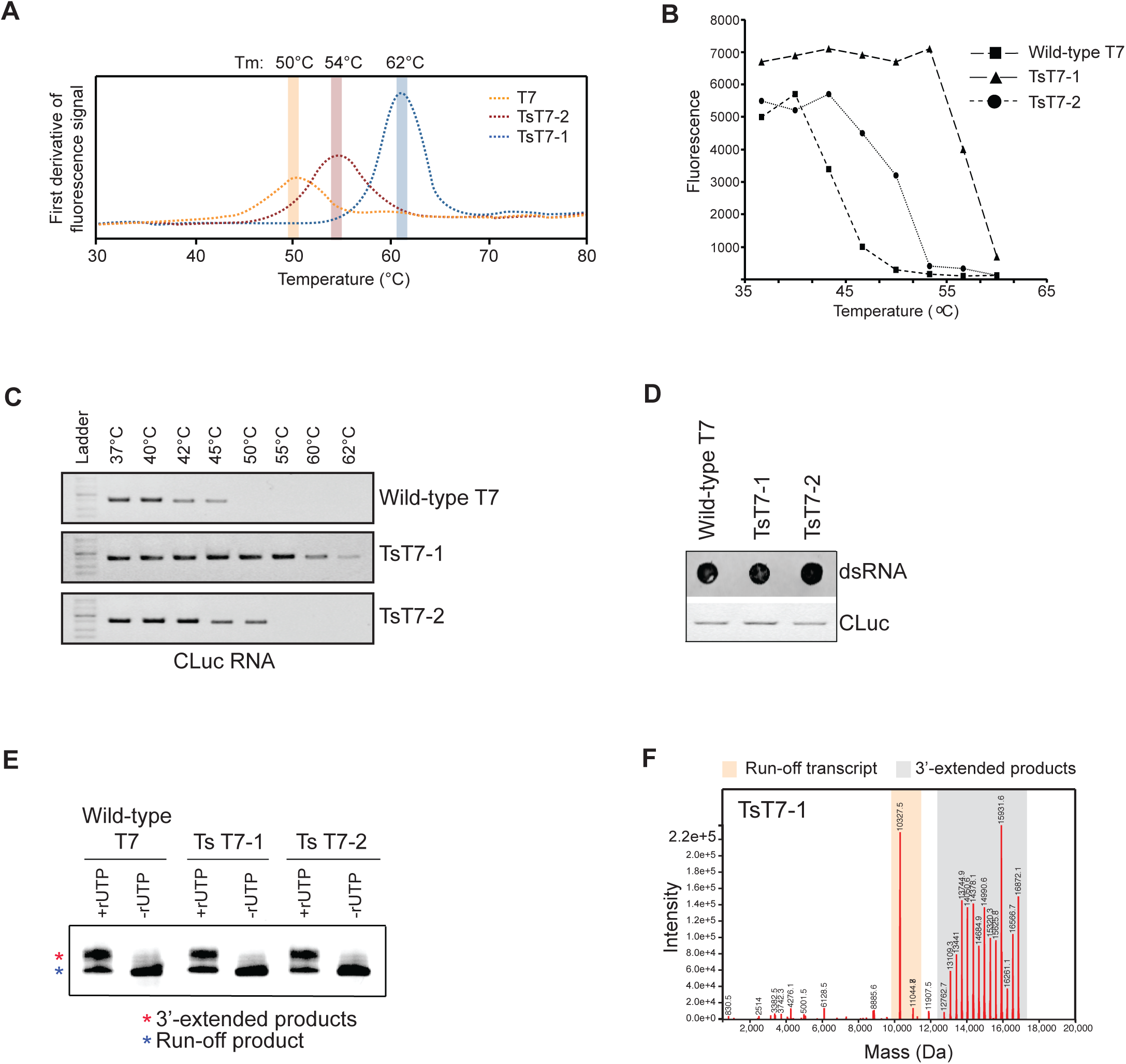
Thermostable (Ts) RNAPs are active at high temperatures and do not affect 3’ extended RNA by-product formation. **A)** Melting temperatures of wild-type T7 RNAP (orange), TsT7-1 (blue) and TsT7-2 (red) determined by nano-differential scanning fluorimetry. **B)** Molecular beacon assay for efficiency of *in vitro* transcription with TsT7-1 or TsT7-2 compared to wild-type RNAP at temperatures ranging from 37°C to 62°C. **C)** Gel electrophoreses analyses of CLuc RNA synthesized with either wild-type RNAP, TsT7-1 or TsT7-2 RNAPs at temperatures ranging from 37°C to 62°C. **D)** dsRNA immunoblot using J2 antibody and gel electrophoresis analysis of CLuc RNA synthesized from IVT reactions performed at 37°C using wild-type T7, TsT7-1 and TsT7-2 RNAPs. **E)** Denaturing gel analysis of IVT reactions using Template IVT short-1 with Ts RNAPs compared to wild-type RNAP. Reactions were performed with either four NTPs (+rUTP) or with three NTPs (-rUTP). **F)** Intact mass spectrometry of short IVT products from TsT7-1 under standard conditions (37°C) with products greater than 30-nucleotide highlighted in grey and the expected run-off transcripts in orange (including n+1, n+2 products from non-templated additions).

### Thermostable RNAPs synthesize 3’-extended dsRNA byproducts

The two thermostable RNAPs tested in this study have different sets of mutations that confer thermostability to the polymerase [25]. In order to determine whether the mutations altered the activity of the enzyme, and in turn the formation of the spurious byproducts, IVT reactions were performed under standard conditions (37°C for 1 hour) and analyzed by either the dsRNA immunoblot assay (for CLuc mRNA) or by intact mass analysis (for Template IVT short 1). The immunoblot assay showed similar levels of dsRNA contaminants in the reactions with both thermostable RNAPs (Figure 2D). Additionally, denaturing gel electrophoresis and intact mass analysis of the short IVT RNA showed the presence of the 3’-extended RNA products (Figures 2E, F), suggesting that the mutations introduced into the thermostable RNAPs do not affect the synthesis of the 3’-extended byproducts of the reaction.

### 3’-extended dsRNA byproducts are reduced in high-temperature IVT reactions

We tested the effect of temperature on the formation of the 3’-extended byproducts by performing the transcription reactions at temperatures ranging from 37°C to 60°C. Analysis of the short IVT template 1 products by denaturing gel electrophoresis showed the presence of the two RNA products (run-off product and 3’-extended product) in IVT reactions performed at 37°C to 48°C using TsT7-1 (Figure 3A). However, for transcription reactions that were performed at temperatures greater than 48°C, a single RNA product corresponding to the run-off transcript was observed, suggesting a reduction of the 3’-extended byproducts without loss of the run-off product (Figure 3A). Even though the IVT products were analyzed in a denaturing gel, we wanted to confirm that the absence of the higher-molecular-weight species was not merely due to a change in the conformation of the RNA at the higher temperature. Therefore, we subjected the IVT reactions (from IVT template short 1 and 2) to intact mass analysis. Analyses of the RNA products synthesized at 50°C (the optimum temperature for TsT7-1) showed presence of the run-off transcript and RNA products with the non-templated additions, but not the 3’-extended products observed in the 37°C reactions (Figure 3B). Indeed, the reaction products at 50°C were similar to those observed in the -rUTP and -rCTP reactions in Figure 1D and 1E, respectively. Similar to TsT7-1, IVT reactions performed at 50°C with TsT7-2 also showed reduced amounts of the 3’-extended products (Supplementary Figure S6). We further tested whether high-temperature transcription could affect the formation of the dsRNAs with templates of different sequences. As expected, when we analyzed the dsRNA contaminants formed in IVT reactions of the CLuc RNA using the dsRNA immunoblot, we observed a reduction in the total dsRNA levels when IVT reactions were performed at 48°C or higher temperatures (Figure 3C). This is consistent with the reduction in the amounts of the 3’-extended RNA products from the shorter templates and suggests that, even though the RNA immunoblot is detecting a pool of dsRNA contaminants, the majority of them are likely 3’-extended products, whose formation is reduced in high-temperature transcription.

**Figure 3.**
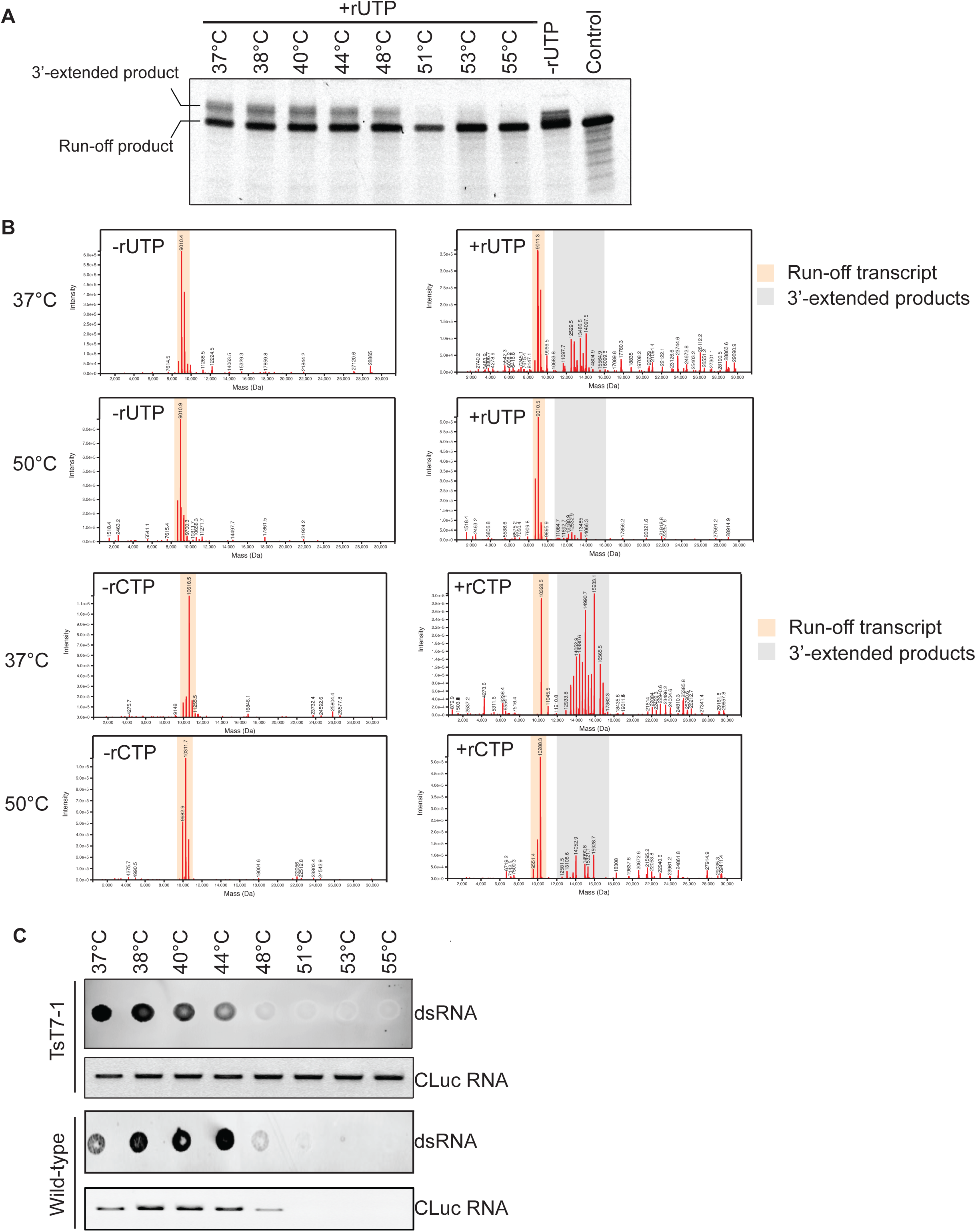
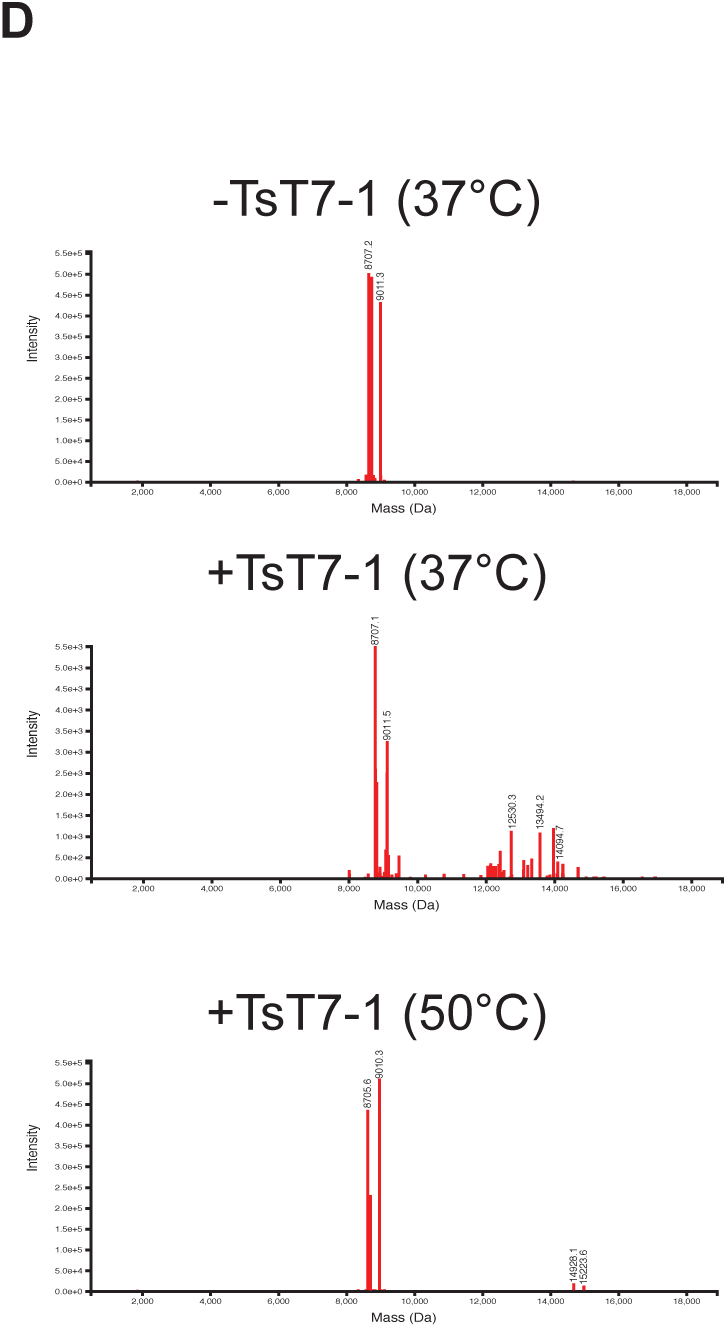
High-temperature *in vitro* transcription with TsT7-1 leads to reduction in 3’-extended byproducts. **A)** Denaturing gel analysis of IVT products synthesized from short IVT template-1 using TsT7-1 performed at a temperature range of 37°C to 55°C. Chemically synthesized short IVT template 1 RNA was used as a control. **B)** Intact mass spectrometry analysis of Template IVT short-1 and Template IVT short-2 using TsT7-1 under standard conditions (all four rNTPs; +rUTP or rCTP) or lacking one rNTP (-rUTP or -rCTP) performed at either 37°C or 50°C. Expected run-off transcript (including n+1 and n+2 products) is highlighted in orange and products larger than 30 nucleotides are highlighted in grey. **C)** dsRNA immunoblot with J2 antibody and gel electrophoresis analysis of CLuc RNA synthesized with TsT7-1 or wild-type RNAP at a temperature range from 37°C to 55°C. **D)** Intact mass spectrometry analysis of IVT reactions in presence or absence of TsT7-1 on chemically synthesized RNA (n, n+1, n+2) performed at either 37°C or 50°C. Products larger than 30 nucleotides are highlighted in grey.

### High-temperature IVT prevents the self-priming of the run-off product

Earlier work has demonstrated that the 3’-extended byproducts are synthesized by self-extension of the run-off transcript, leading to synthesis of considerably longer RNA products during IVT [11, 13, 20]. Therefore, we tested whether the 3’-extended byproducts we observed from our templates were synthesized by the same mechanism and whether high-temperature IVT prevented the self-extension of the run-off product by T7 RNAP. We chemically synthesized a 30-nt RNA (corresponding to IVT Template short-1) and incubated it in IVT reactions with NTPs (5 mM each) and TsT7-1 RNAP, but in the absence of any T7-promoter-containing DNA template. For reactions that were performed at 37°C, intact MS analysis of the RNA products showed presence of the 3’-extended products but at reduced levels as compared to when the reaction was done with the DNA template (Figure 3D). This suggests that in absence of a promoter sequence, the T7 RNA polymerase can bind an RNA and cause RNA-dependent RNA extension. In contrast, reactions performed at 50°C did not show any detectable 3’-extended products (Figure 3D). In conclusion we find that consistent with previous studies, T7 RNA polymerase can efficiently 3’-extend run-off transcripts [20], and that high-temperature IVT reduces the 3’-extension of the run-off product.

### High-temperature IVT does not affect formation of antisense RNA byproducts

A previous report has demonstrated the synthesis of antisense RNA molecules that can base-pair with the run-off transcript and contribute to formation of dsRNA contaminants in the IVT reaction [21]. We wanted to investigate whether high-temperature IVT affected formation of antisense-dependent dsRNA structures. Because we were unable to detect the formation of the antisense RNA in any of our templates under native conditions (data not shown), we tested the same sequence that was reported in the previous study (template 512B)[21]. With the 512B template, we were able to detect the formation of RNA molecules that were sensitive to RNase III treatment when transcription was performed with either wild-type RNAP (Figure 4A) or the TsT7-1 RNAP (Figure 4C). The RNase III-sensitive species were only detected under native gel electrophoresis conditions and not under denaturing gel electrophoresis conditions, suggesting that the run-off transcript and the antisense products had similar migration patterns and were not the same as the 3’-extended products that we and others [20] have observed from other templates. Furthermore, the immunoblot assay was able to detect dsRNA-specific signals from the IVT reactions using the 512B template (Figure 4B), suggesting that dsRNA species were formed and that the J2 antibody could recognize them irrespective of their nature. To investigate the effect of high-temperature transcription on the formation of the antisense dsRNA species, IVT was performed with TsT7-1 at either 37°C or 50°C. Interestingly, TsT7-1 did not have an effect on the formation of the RNase III-sensitive antisense-mediated dsRNA species at higher temperatures (Figure 4C). This suggests that the mechanism by which the antisense-mediated dsRNAs are formed is different than that for the 3’-extended dsRNA products, and the conditions that could affect the formation of these are also distinct. Lowering the concentration of magnesium ions in the reaction has been suggested to affect the antisense:sense dsRNA formation [21]. We tested the effect of lowering the magnesium concentration on formation of dsRNA byproducts. As expected, overall RNA yield was reduced rather than just that of the byproducts (data not shown) [34–36].

**Figure 4.**
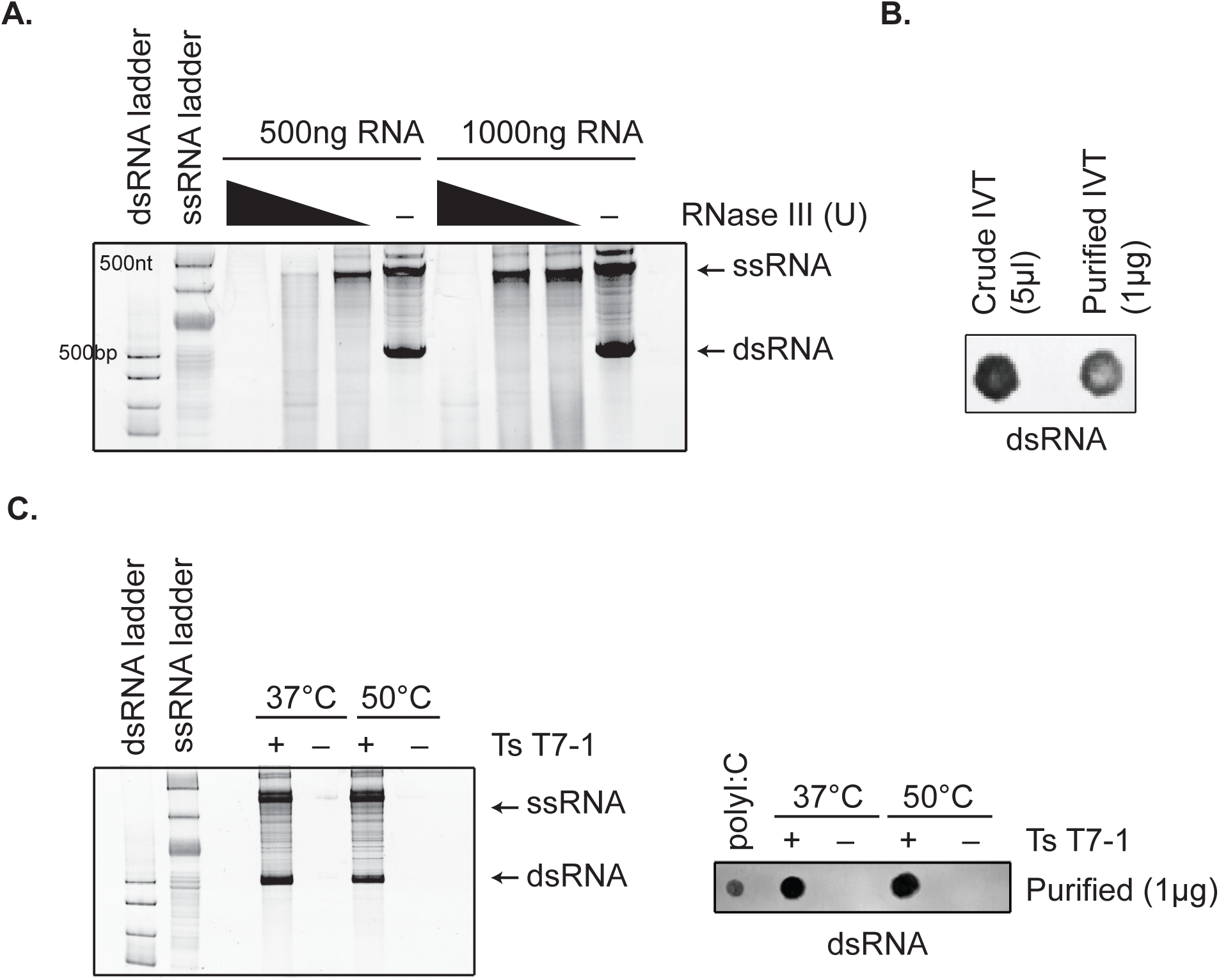
High-temperature *in vitro* transcription does not affect antisense dsRNA by-product formation. **A)** Native gel electrophoresis analyses of IVT reactions on 512B DNA template using wild-type T7 (37°C) with/without RNase III treatment. **B)** dsRNA immunoblot with J2 antibody on IVT reactions (crude and purified) with 512B template. **C)** Native gel electrophoresis analyses and dsRNA immunoblot analysis of 512B IVT reactions conducted with TsT7-1 at 37°C *vs*. 50°C.

### Antisense byproduct formation is sequence specific

The immunoblot assay showed reduction in the dsRNA content when transcription was performed at high temperature for multiple templates (Supplementary Figure S7). Furthermore, the immunoblot could detect dsRNA irrespective of its source or nature (Figure 1A, Figure 4B). The fact that high-temperature transcription reduced dsRNA signals from multiple templates (with the exception of the 512B template) suggested that the majority of the dsRNA byproducts were 3’-extended and that the antisense byproducts from the 512B template might be sequence specific. To understand the formation of the antisense RNA byproducts and the effect of high temperature on their synthesis, we constructed chimeric templates in which the 3’-end sequence of the 512B template was altered. Interestingly, moving the terminal 25 base pair segment away from the 3’ end of the template towards the 5’-end, reduced the level of the antisense dsRNA products from the 512B template when IVT was performed with either wild-type T7 RNAP (Figure 5A) or TsT7-1 RNAP (Figure 5B) under standard reaction conditions. We hypothesized that since the presence of such a sequence at the 3’-end of the template increased the propensity of the polymerase to switch to the non-template DNA strand and initiate transcription, then adding sequences from the CLuc mRNA 3’-end (512B::CLuc) would alleviate this problem. As expected, when we analyzed formation of the antisense dsRNA using the 512B::CLuc template, we observed a reduction in the formation of the antisense dsRNA in the reaction (Figure 5C). Taken together, these data demonstrate that formation of the antisense RNA in the 512B template requires a specific sequence to be present at the 3’-end of the DNA template. Furthermore, it shows that the CLuc RNA 3’ sequence does not have a similar sequence element and, therefore, explains the observed reduction in dsRNA byproducts in this mRNA preparation after high-temperature IVT (Figure 3C).

**Figure 5.**
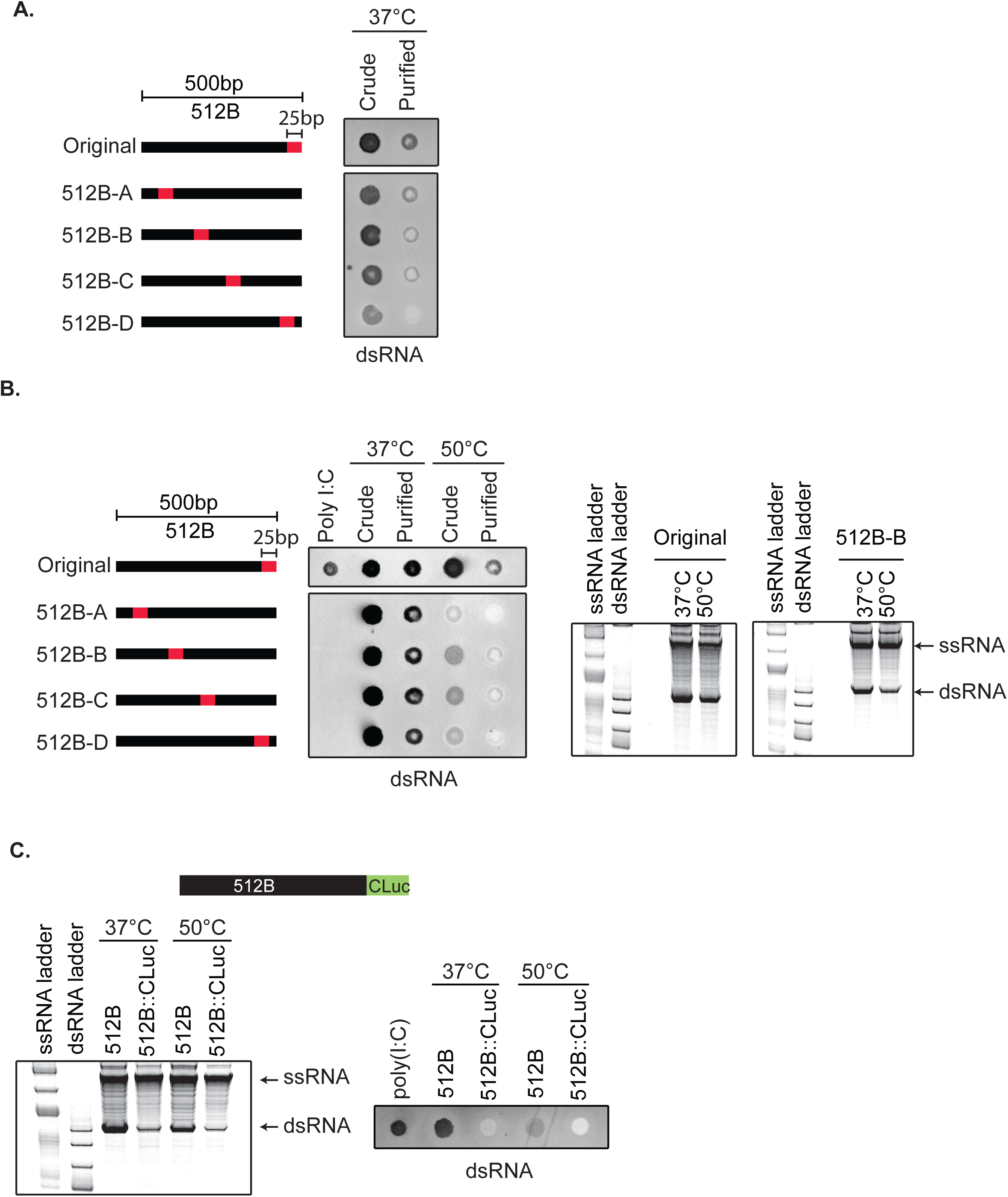
Antisense dsRNA by-product formation is template 3’ sequence dependent. **A)** dsRNA immunoblot using J2 antibody on IVT reactions with wild-type T7 at 37°C performed on modified 512B templates in which the 3’-terminal 25 bp sequence was moved to various positions within 512B template (512B-A to 512B-D). **B)** dsRNA immunoblot and native gel analysis of modified 512B templates (512B-A to 512B-D) using TsT7-1 at 37°C vs. 50°C. **C)** A chimeric template was generated in which 26 bp of the CLuc 3’-end sequence was added to the 3’-end of 512B template (denoted 512B::CLuc). Native gel electrophoresis analyses and dsRNA immunoblot analysis of IVT reactions of the 512B::CLuc chimeric template compared to the original unmodified 512B template. Reactions were performed with TsT7-1 at either 37°C or 50°C for one hour.

We next tested the effect of high-temperature transcription on dsRNA by-product formation using the chimeric templates where the extent of antisense RNA product formation has been altered. We observed a reduction in the total amount of dsRNA by immunoblot, as well as a reduction in the dsRNA products from the sense:antisense pairing at higher temperature with TsT7-1 (Figure 5B). Finally, by truncating the 3’ end of the 512B template by 50 bp or 200 bp to eliminate the sequence-specific effect of the template on formation of the antisense RNA by-product, we could detect reduction in dsRNA levels at higher temperatures (Supplemental Figure S8).

### Presence of a poly-T template sequence affects formation of the antisense but not the 3’-extended byproducts

The mechanistic insights on the formation of the two kinds of dsRNA byproducts have come from short RNA substrates. However, for the synthesis of functional mRNAs, a poly-A tail is required, and for most applications, it is encoded from the DNA template. We investigated whether the presence of a poly-A tail in the 3’ end of the run-off transcript (poly-T tract in the template DNA) affected formation of the undesired dsRNA byproducts. We generated CLuc DNA templates with varying lengths (30, 60, 120 base pairs) of a poly-T sequence at the 3’-end, performed IVT reactions under standard conditions (37°C for 1 hour), and analyzed the presence of the dsRNA byproducts with the immunoblot assay. No substantial differences in the formation of dsRNA byproducts was observed when templates with varying lengths of poly-T were used (Figure 6A). This observation suggests that at least one, if not both types of the dsRNA byproducts were still synthesized when poly-T sequences are present at the 3’ end of the DNA template.

**Figure 6.**
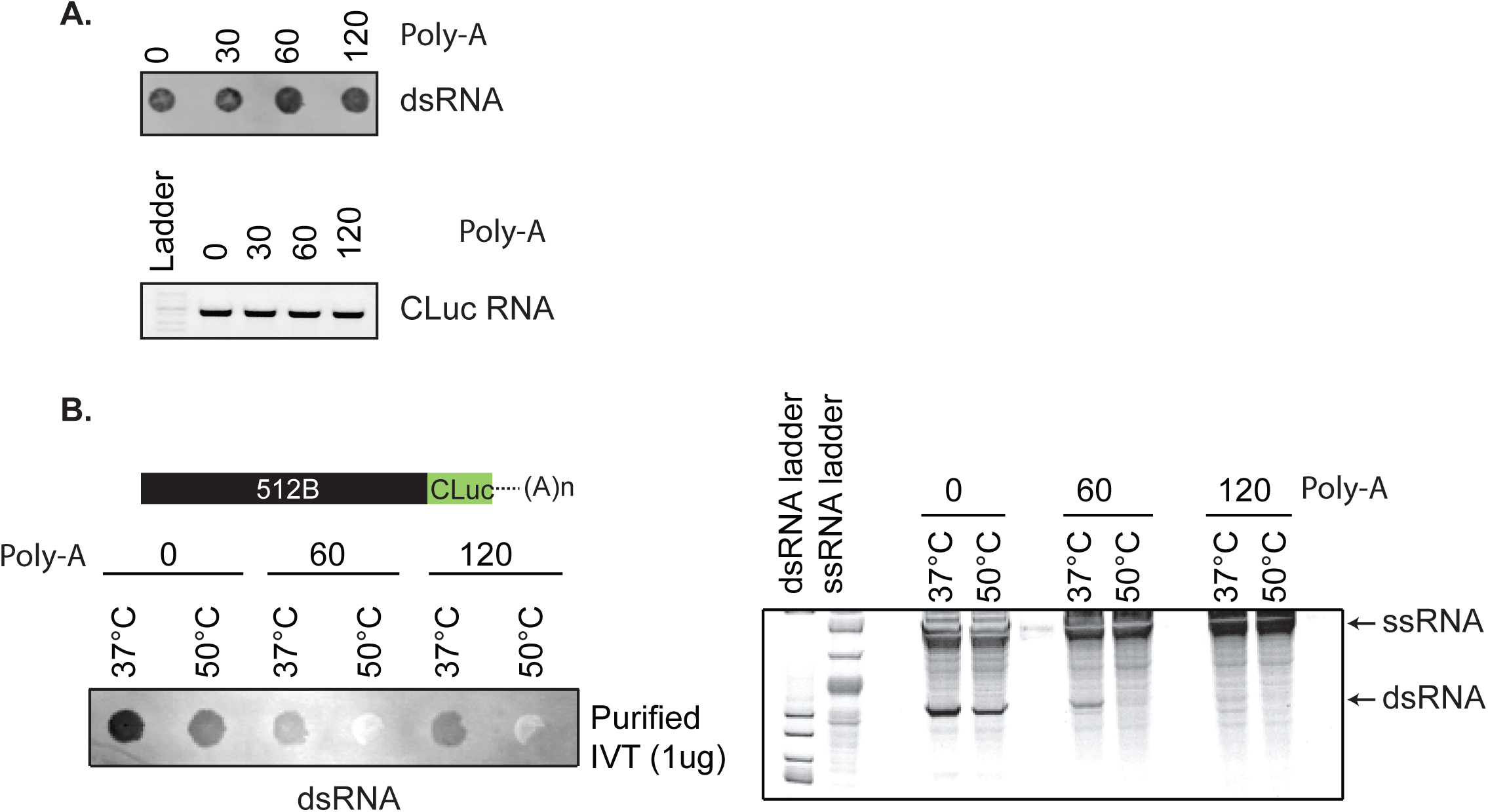
Template-encoded Poly(A) tailing reduces antisense by-product formation. **A)** dsRNA immunoblot with J2 antibody and gel electrophoresis analysis of CLuc RNA synthesized from CLuc templates with varying length (30 bp, 60 bp, 120 bp) of poly-T sequence at 3’-end under standard conditions (5 mM rNTPs, 37°C for 1 hour). **B)** Immunoblot and native gel electrophoresis analysis of IVT reactions on 512B::CLuc chimeric template with poly-T (60 bp and 120 bp) sequence at the 3’-end. IVT reactions were performed at 37°C or 50°C.

Based on our data from the 512B-chimeric templates (Figure 5A-C), we hypothesized that presence of a poly-T sequence in the template might be able to override the effect of the antisense-promoting sequence. Varying lengths (60 and 120 base pairs) of poly-T sequences were added to the 3’ end of the 512B sequence to test this hypothesis (Figure 6B). Analyses of the RNA products from the poly-T-containing 512B templates showed a reduction in the antisense dsRNA species, suggesting that the effect of sequence elements that drive the T7 RNAP towards switching to the non-template strand can, in part, be mitigated by the addition of the poly-T sequences (Figure 6B). To test whether addition of the poly-A tail affected the formation of the 3’-extended dsRNA products, we added poly-T sequence to the short RNA templates (Template IVT short 1). The synthesis of the 3’-extended products was not affected when poly-T sequences were present (Supplemental Figure S9). This suggests that even in the presence of a poly-A sequence at the 3’-end of the run-off transcript, the RNAP can most likely re-bind the run-off product and undergo efficient self-priming.

### Reduced immune response from functional mRNAs synthesized at higher temperatures

Because presence of dsRNAs in the IVT reaction can stimulate a cellular immune response, we asked whether the reduction in dsRNA byproducts observed after high-temperature IVT was sufficient to reduce the immune response observed upon introduction of synthetic mRNAs in cells. We analyzed the immune response from capped and poly-A tailed CLuc mRNAs synthesized with either wild-type T7 or TsT7-1 RNAPs. Furthermore, to ensure that we are measuring the immune response exclusively from the dsRNA byproducts, we introduced pseudouridine modifications in the mRNAs to evade immune response from TLR7 and TLR8 [7, 37]. Comparison of luciferase activity from HEK293 cells that were transfected with CLuc mRNAs synthesized with either wild-type T7 or TsT7 RNAP showed no considerable differences in expression, suggesting that mRNAs synthesized with TsT7-1 at 50°C were functional (Figure 7A). Comparison of the IFN-α levels from monocyte-derived dendritic cells transfected with CLuc mRNA (transcribed at 37°C or 50°C with wild-type T7 or TsT7-1 respectively) showed that crude IVT CLuc mRNA (without extensive purification) synthesized with wild-type T7 RNAP at 37°C results in secretion of high levels of IFN-α as compared to that seen with control total rat RNA (Figure 7C, D). As a comparison for efficient removal of dsRNA byproducts from the mRNA preparation, CLuc mRNAs synthesized at 37°C were subjected to high-performance liquid chromatography (HPLC) purification, and fractions either devoid of the dsRNA by-products (Fraction II;) or enriched (Fraction I and Fraction III) were transfected. Fraction II resulted in reduced secretion of IFN-α (Figure 7B). Interestingly, CLuc mRNA synthesized with TsT7-1 at 50°C followed by a silica column cleanup (but no HPLC purification) also showed reduced IFN-α levels as compared with that seen for RNA synthesized at 37°C. The IFN-α levels from the high-temperature IVT mRNAs were comparable to the HPLC-purified Fraction II, suggesting that part of the immune response generated from the presence of the 3’-extended dsRNA byproducts in the reaction can be overcome by high-temperature IVT.

**Figure 7.**
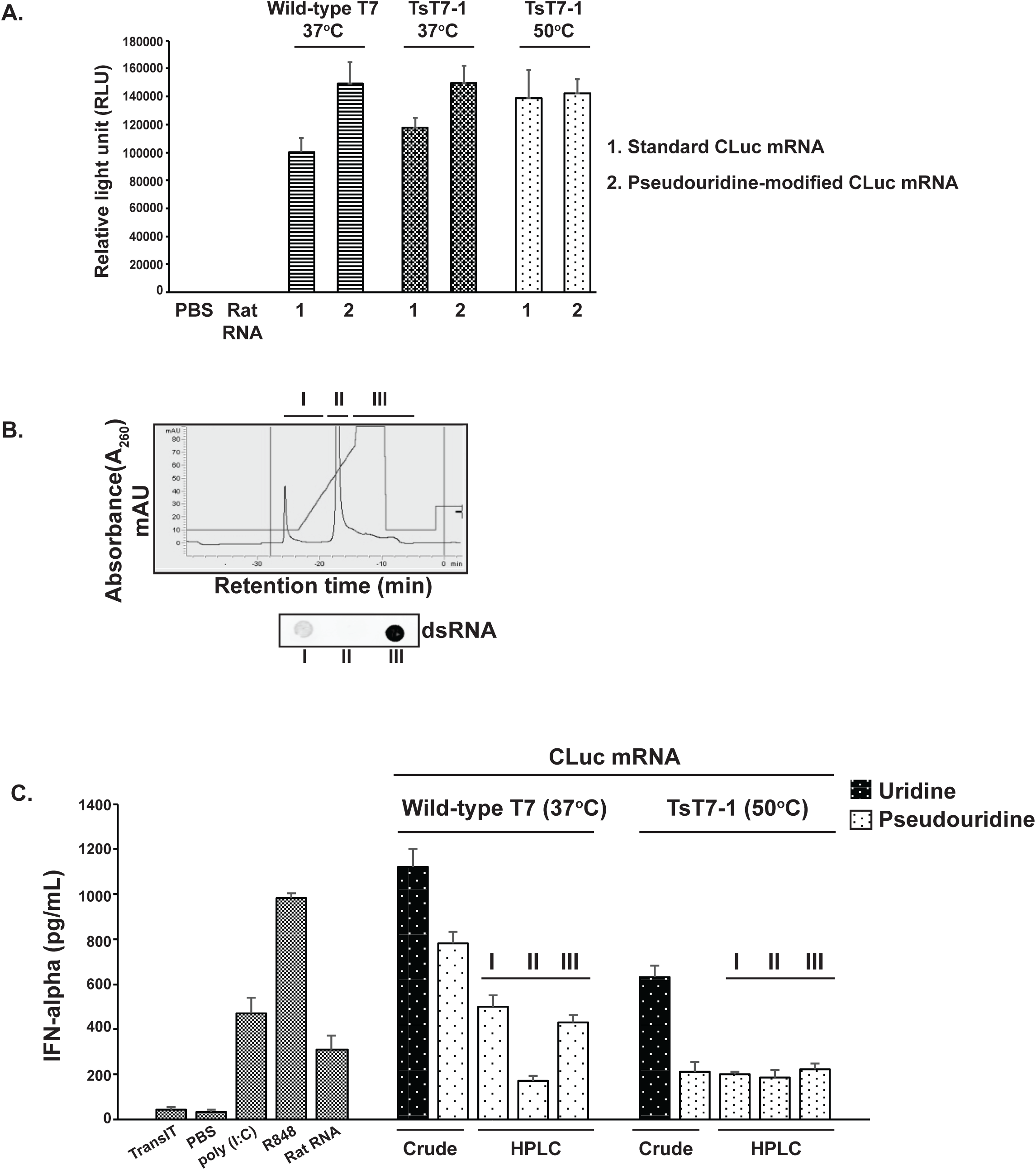
mRNAs generated by high-temperature *in vitro* transcription with TsT7-1 are functional and have reduced immunogenicity *in vivo*. **A)** Luciferase activity from HEK293 cells transfected with CLuc mRNA (unmodified or modified with pseudouridine) synthesized with wild-type T7 or TsT7-1 RNAP at 37°C or 50°C. **B)** Representative chromatogram for CLuc mRNA separated on an HPLC column. Absorbance at 260nm representing the RNA products in the IVT reaction. dsRNA immunoblot with J2 antibody on fractions collected from the HPLC purification. **C)** IFN-α levels in supernatant collected from DCs transfected with either crude IVT CLuc mRNA or CLuc mRNA purified with HPLC. IFN-α levels in the supernatant were measured 24 hours post-transfection. Error bars are standard error of the mean. Poly(I:C), a synthetic analog of dsRNA and Resiquimod (R848), an activator of Toll-like receptors were used as controls for interferon activation.

## DISCUSSION

It is well established that *in vitro* transcription results in formation of dsRNA byproducts in the reaction, which can have immunostimulatory effects when introduced *in vivo* [5, 10, 11]. However, the precise nature and source of these byproducts is neither well understood nor well characterized. During the course of this work, two independent studies aimed towards understanding the nature of these dsRNA byproducts were published [20, 21]. Interestingly, the nature of the dsRNA byproducts described in these two studies were very different. The discrepancies between these two studies have not been systematically investigated and reconciled. Furthermore, it is not clear whether synthesis of the 3’-extended products and the antisense byproducts are dependent on each other and whether the parameters that affect the formation of one type of by-product can alter the synthesis of the other type.

Here we demonstrated that high-temperature IVT using thermostable T7 RNAP reduces the formation of dsRNA byproducts. Our analyses of the dsRNA byproducts by using multiple approaches demonstrated that the predominant spurious product generated from the majority of the templates was a 3’-extended species of the main run-off transcript that was able to form RNase III-sensitive dsRNA structures (Figure 1B, 1C, 3D). This is consistent with observations from the study by Gholamalipour *et al*. that also identified the 3’-extended products as the predominant type of by-product in IVT reactions and identified the run-off products to be the templates for synthesis of these spurious species [20, 22]. The mechanism for formation of these 3’-extended products has been suggested to be the re-binding of the RNA polymerase to the run-off product when the RNA products have accumulated in sufficient amounts in the reaction. The re-binding of the RNA polymerase could essentially result in the folding back of the RNA onto itself followed by extension using the same molecule as the template (self-extension). Our characterization of RNA produced by IVT at higher temperatures demonstrated a reduction in the synthesis of the 3’-extended byproducts when the reactions were performed at temperatures greater than 48°C (Figure 3A-C). It is tempting to speculate that higher temperature alters either the re-binding of the RNAP to the run-off transcript or the folding back and efficient self-priming of the transcript.

In some cases, as exemplified by a specific DNA template in both previous [21] and our studies, formation of antisense transcript with a promoter-independent transcription initiation mechanism has been demonstrated where the dsRNA is formed by hybridization of the run-off transcript with the antisense transcript [21]. Our analyses of this specific DNA template (Figure 5A-C) demonstrate that formation of these byproducts is sequence specific and requires that sequence to be present at the 3’-end of the template DNA. The exact mechanism by which this sequence induces the transcription of the non-template strand by T7 RNAP needs to be investigated.

Our observations that high-temperature IVT reduced the amount of 3’-extended products, but had little to no effect on the synthesis of the antisense RNA by-product (Figure 4C) suggests that two competing mechanisms are at play toward the end of run-off transcription process. In one such mechanism, as suggested by our data (Figure 5A) and others [20], the polymerase extends the run-off product to form the 3’-extended dsRNA byproducts. The self-extension of the runoff transcript is reduced during high-temperature IVT. However, in the presence of a sequence at the 3’ end of DNA template that promotes the RNAP toward non-template strand switching, the RNAP initiates transcription on the non-template strand, which results in the formation of the antisense product (Figure 4A). Altering the reaction temperature has no effect on this process (Figure 4C). In presence of a sequence that promotes transcription from the non-template DNA strand, 3’-extension might not occur at high enough levels because, instead of re-binding the run-off transcript, the polymerase initiates transcription from the non-template strand. The effects of high temperature on the IVT process were apparent in the 512B template when the 3’ DNA template sequence was either moved (Figure 5D) or substituted with the 3’ sequence of CLuc DNA (Figure 3C, 5C), which suggests that in the absence of the sequence that promotes synthesis of the antisense RNA byproduct, 3’-extended byproduct formation becomes more pronounced. This is reinforced by the observation that we could not detect 3’-extended products from the 512B template (data not shown). We hypothesize that re-binding of the RNAP to the run-off transcript might be overridden by template switching of the RNAP. The synthesis of the 3’-extended product has been shown to be dependent on the accumulation of the run-off transcripts [20]. Martin and colleagues have hypothesized that during early IVT, the RNAP binds the promoter and generates run-off products through the canonical “on pathway”. However, when the run-off transcript accumulates to a certain level, the RNAP participates in a competing “off pathway” wherein it re-binds the run-off RNA, which results in self-primed extension. Based on our data, we propose that presence of a sequence at the 3’ end of the template that allows the RNAP to initiate transcription of the antisense product drives the RNAP to a DNA-sequence-dependent “pseudo-on pathway” that results in reduction of 3’-extension (Figure 8).

**Figure 8.**
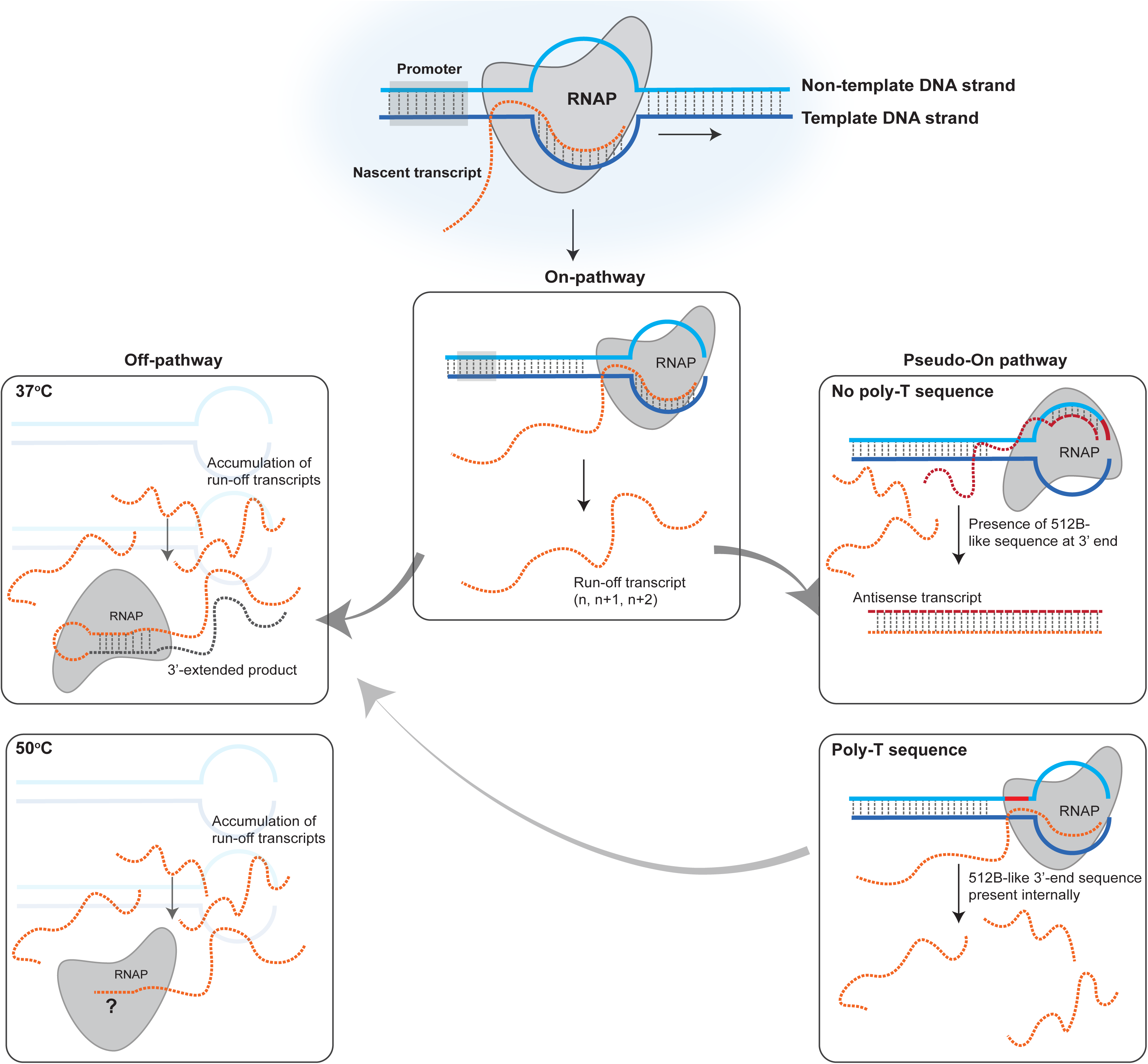
Schematic illustration of competing pathways resulting in formation of dsRNA byproducts during *in vitro* transcription. The canonical “on-pathway” results in promoter-dependent transcription initiation and synthesis of the run-off products [20]. Accumulation of the run-off transcripts in the reaction result in a competing “off-pathway” wherein the RNAP re-binds the run-off product and results in self-extension of the run-off transcript. At temperatures higher than 48°C, the 3’-extension of the run-off products are reduced (due to either reduced re-binding of the RNAP to the run-off product or the folding of the run-off product). In presence of a sequence at the 3’ end of the template (similar to the 512-B 3’-end sequence), the RNAP initiates transcription of the antisense RNA product in a “pseudo-on pathway”. Presence of poly-T sequences prevent the formation of the antisense RNA products. Presence of a poly-A sequence at the 3’-end of the run-off transcript does not affect the “off-pathway” at 37°C and can lead to synthesis of longer products if reactions are performed under standard conditions.

Functional mRNAs require a poly-A tail for stability and efficient protein expression. Poly-A tails are added to the synthetic RNA either post-transcriptionally with poly-A polymerase or during transcription via a template-encoded poly-T sequence. Most previous studies have been conducted on short synthetic RNAs that lack a poly-A tail; therefore, not much is known about the effects of a poly-T tract at the 3’ end of the template and its effect on the formation of the spurious byproducts. Our data demonstrates that formation of antisense RNA byproducts, which requires a specific sequence at the 3’-end of the transcript, is affected by the presence of the poly-T sequence (Figure 5C, 6B). Interestingly, formation of the 3’-extended products was not affected by the presence of template-encoded poly-A tails (Figure 6). Other mechanisms for the synthesis of antisense RNA byproducts have recently been proposed [38]; whether or not high-temperature IVT affects the synthesis of these byproducts needs to be investigated.

Recently several different approaches have been proposed to overcome the effects of the dsRNA byproducts *in vivo*. A common approach for removing double-stranded byproducts is to use a chromatography-based purification step after completion of the synthesis reaction [22, 23]. However, implementing such a purification method depending on the nature of the dsRNA byproduct, might not be able to remove all contaminants. Selective binding to cellulose has been suggested [24], but it is not clear whether such a method can distinguish the effects of the intrinsic secondary structures of the RNA. Annealing a DNA oligonucleotide to the 3’ end of the RNA can help overcome generation of some of the dsRNA byproducts [39]; for therapeutic applications, however, such a strategy would require specific removal of the DNA oligonucleotide. Finally, lowering the magnesium ion levels in the *in vitro* transcription reaction has been suggested to reduce levels of the dsRNA byproducts [21], but it also affects the overall yield of the reaction. Furthermore, it is not clear whether the approaches mentioned above will work for the reduction/removal of both kinds of dsRNA byproducts that can be made in the reaction.

Our characterization of the thermostable RNAPs (Figure 2A-B, Figure 7) demonstrate that the RNAs synthesized with the thermostable RNAPs are functional *in vivo* and demonstrate a reduced immune response in dendritic cells due to reduction in 3’-extended byproducts. Furthermore, combination of template-encoded poly-A tailing and high-temperature IVT reduces formation of two types of dsRNA byproducts, improves the purity of RNA, and could potentially alleviate the need for extensive post-synthesis purification.

## SUMMARY

Post-synthesis purification strategies have been shown to successfully eliminate the dsRNA byproducts from the final mRNA preparations, but the cost associated with them, as well as the incompatibility with scale-up processes, makes them impractical for large scale drug development applications. As outlined in this study, high-temperature IVT with thermostable RNAPs provides a novel, simple, and scalable method to synthesize RNA *in vitro,* especially for applications where generating the run-off products without any other byproduct is critical. The generality of this approach for synthesis of mRNA of any sequence makes it easily adoptable and amenable to be integrated into current RNA synthesis workflows.

## ACKNOWLEDGEMENTS

The authors would like to thank Jennifer Ong and Vladimir Potapov for critical discussions and expertise with thermostable RNA polymerases; Jennifer Ong and Tom Evans for comments on the manuscript.

## AUTHOR CONTRIBUTION

B.R. and M.Z.W. conceived and designed the experiments. M.Z.W., B.R., H.A., and G.T. carried out the experiments, B.R. and M.Z.W. wrote the manuscript with input from all authors.

---

**Supplementary Figure S1.**
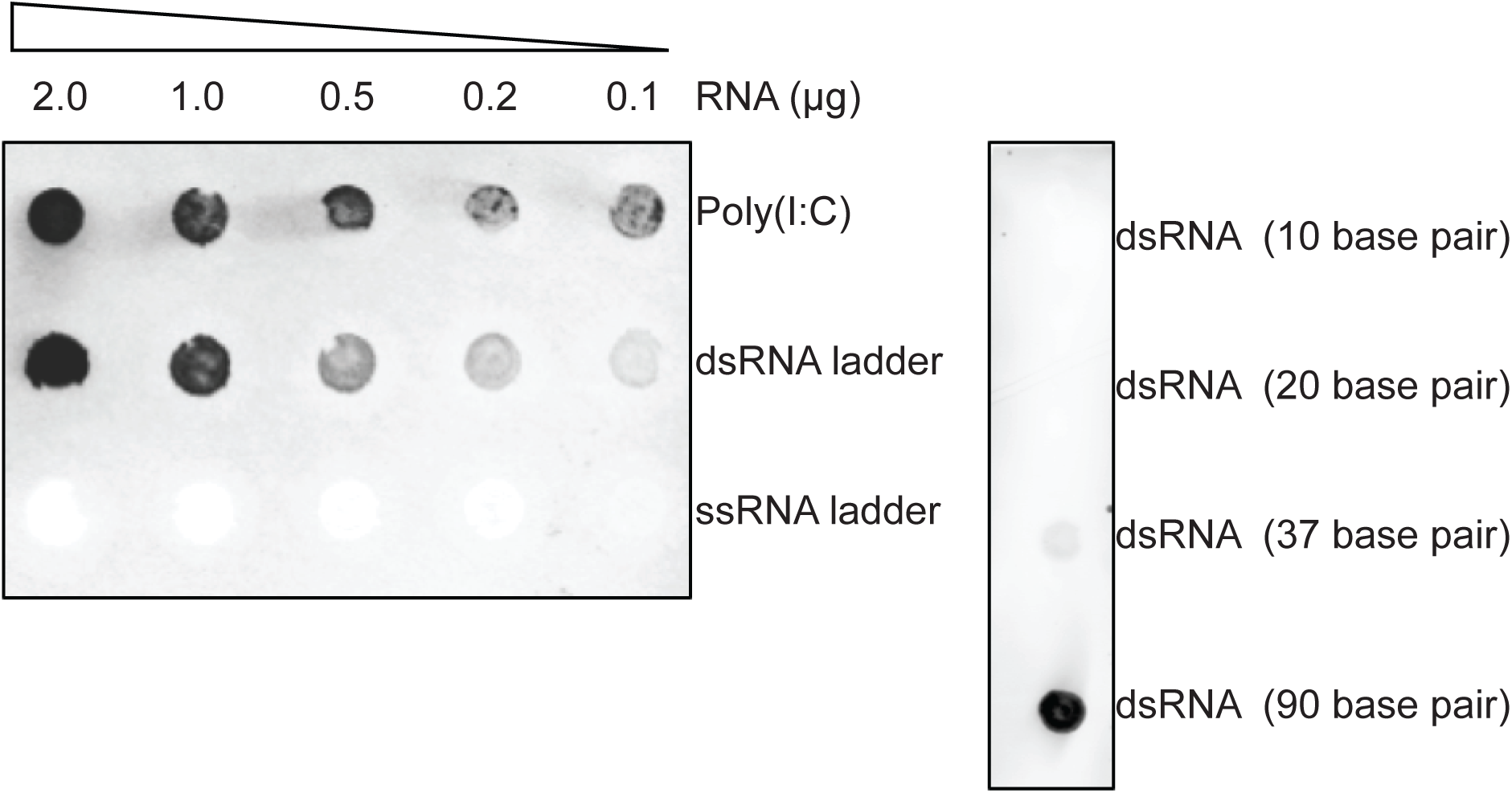
Specificity of anti-dsRNA antibody J2. Immunoblot on synthetic dsRNA of varying sizes (500ng); synthetic analog of dsRNA – poly(I:C) (Invivogen); dsRNA ladder (New England Biolabs); ssRNA ladder (New England Biolabs) with J2 antibody (1:5000; Scicons).

**Supplementary Figure S2.**
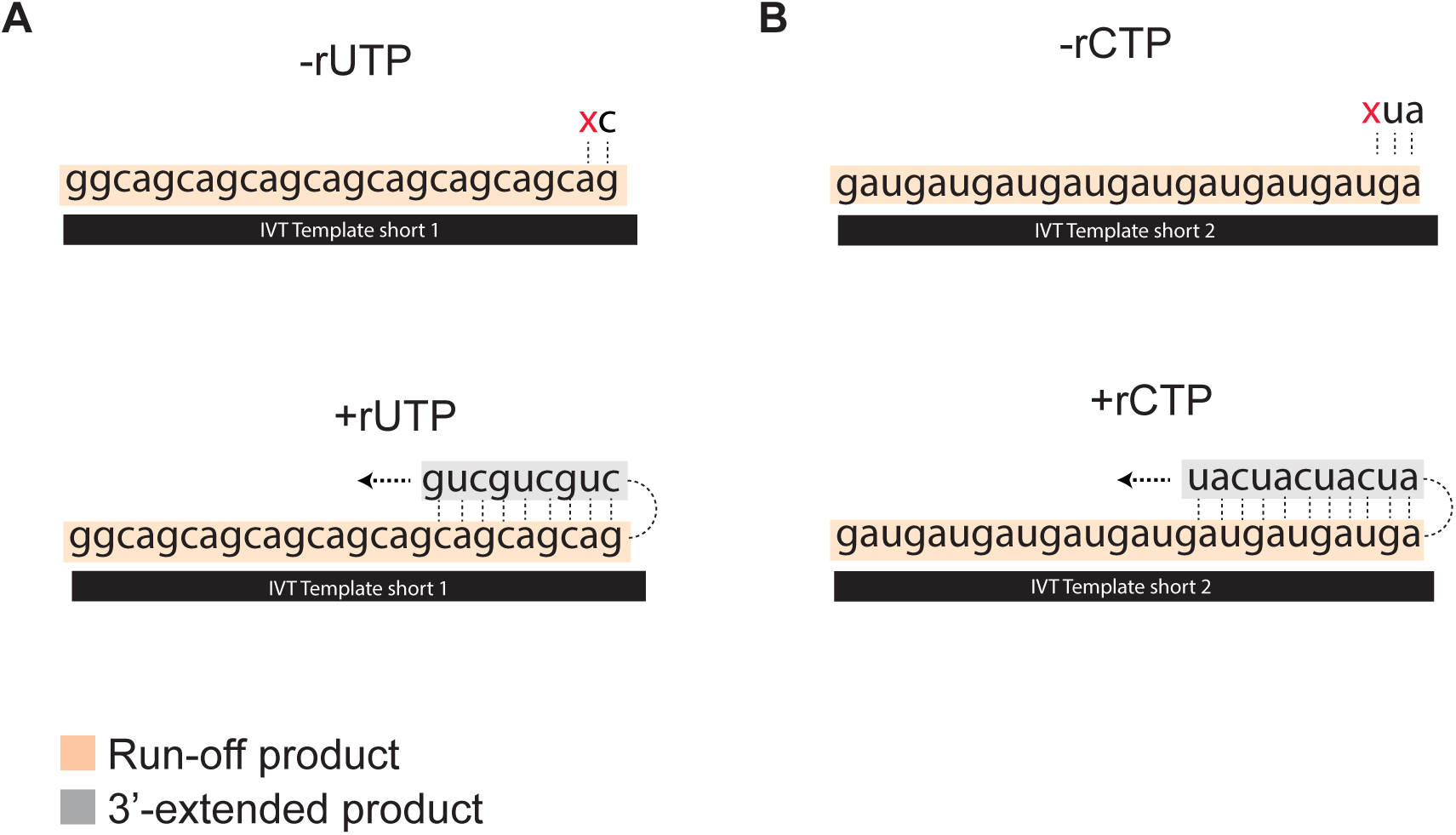
Schematic illustration showing the formation of dsRNA by-products in the presence of all four nucleotides in the reaction for IVT Template short 1 (A) and IVT template short-2 (B) and the formation of run-off product with three NTPs (-rUTP for IVT Template short 1 and –rCTP for IVT Template short 2).

**Supplementary Figure S3.**
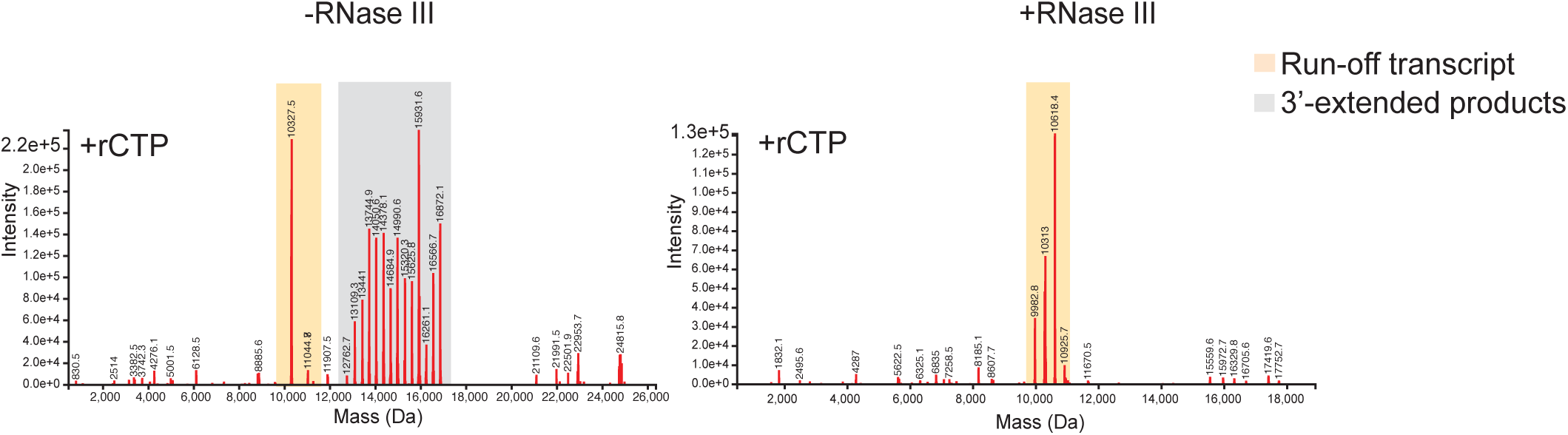
Intact mass spectrometry analyses of *in vitro* transcription reaction with IVT Template short-2 after RNAse III treatment demonstrating reduction in the extended products. *In vitro* transcription reactions were performed with wild-type T7 RNA polymerase at 37°C for 1 hour followed by RNase III treatment. RNase III treated samples were then subjected to intact mass spectrometry analyses.

**Supplementary Figure S4.**
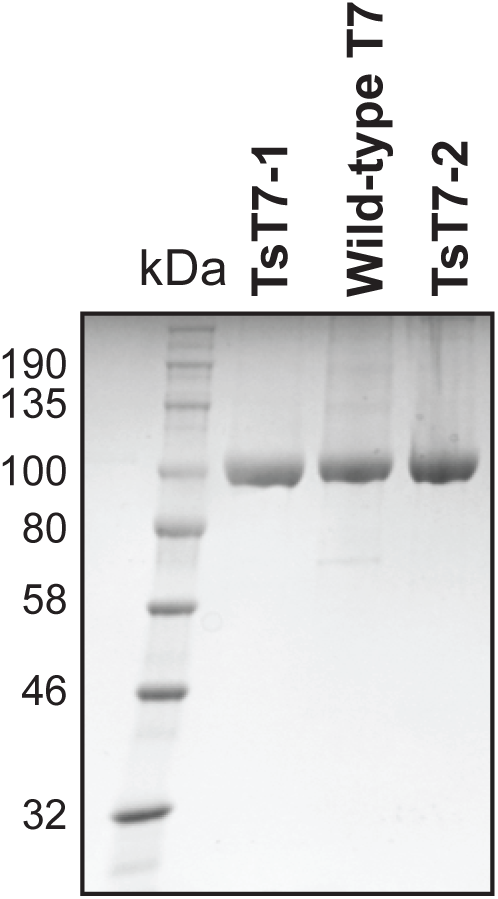
Coomassie-stained SDS polyacrylamide gel showing purity of the RNA polymerase (RNAP) preparations. Equal amounts of protein (wild-type T7 RNAP and TsT7-1 and TsT7-2) were analyzed on an SDS-polyacrylamide gel followed by Coomassie staining.

**Supplementary Figure S5.**
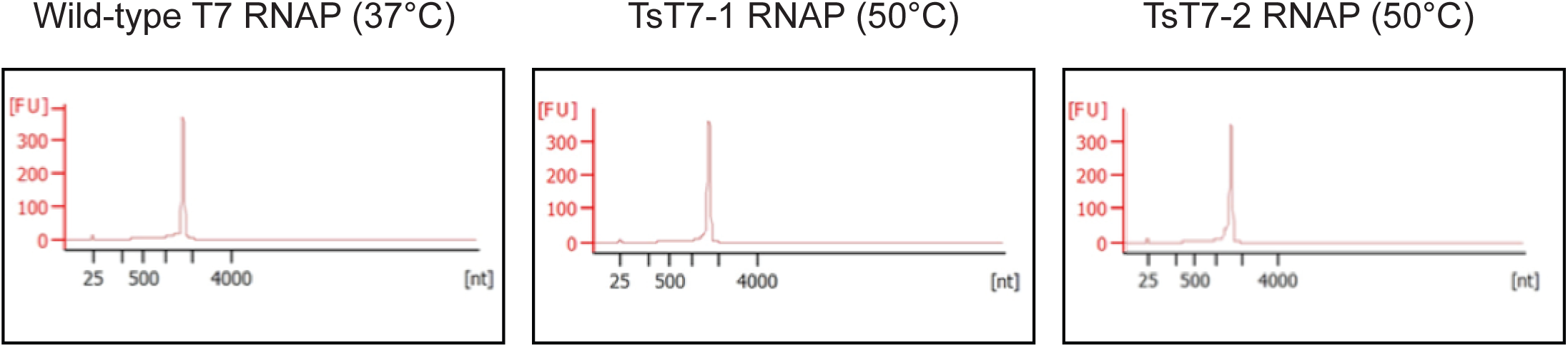
Bioanalyzer trace demonstrating integrity of the CLuc RNA when *in vitro* transcription was performed at 50°C with TsT7-1 and TsT7-2. *In vitro* transcription reactions were performed with either wild-type T7 at 37°C or TsT7-1 and TsT7-2 at 50°C followed by column purification (MEGAclear™ Transcription Clean-Up Kit) and analysis on Agilent 2100 Bioanalyzer.

**Supplementary Figure S6.**
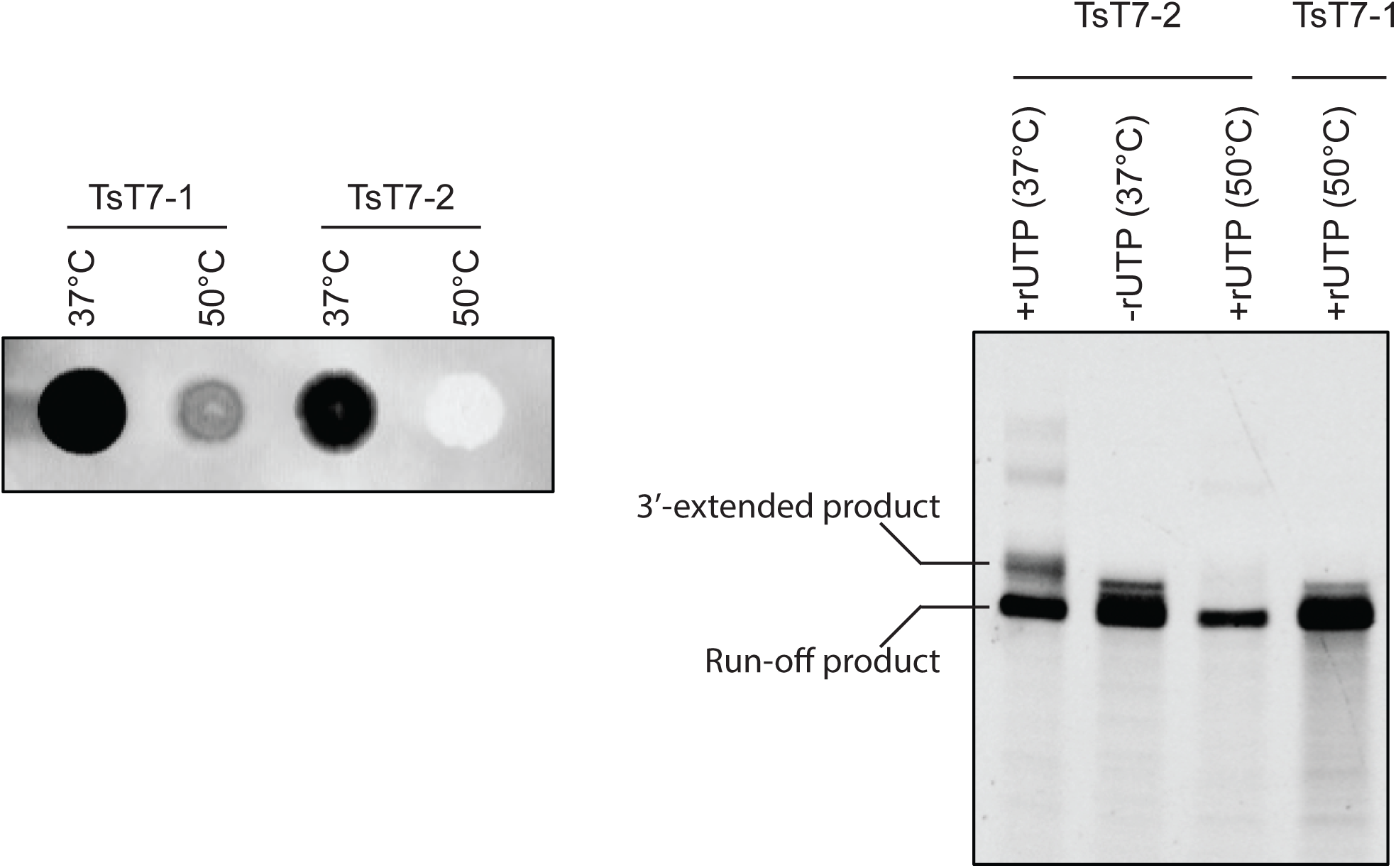
dsRNA immunoblot showing reduction of 3’-extended dsRNA by-products with TsT7-2. Immunoblot using dsRNA antibody J2 on *in vitro* transcription reactions with CLuc template using TsT7-2 at 37°C or 55°C for 1 hour. *In vitro* transcription reactions using short template (30 bp) ran on denaturing gels under standard conditions (four rNTPs), and under conditions where one rNTP is omitted to prevent formation of the dsRNA products (-rUTP).

**Supplementary Figure S7.**
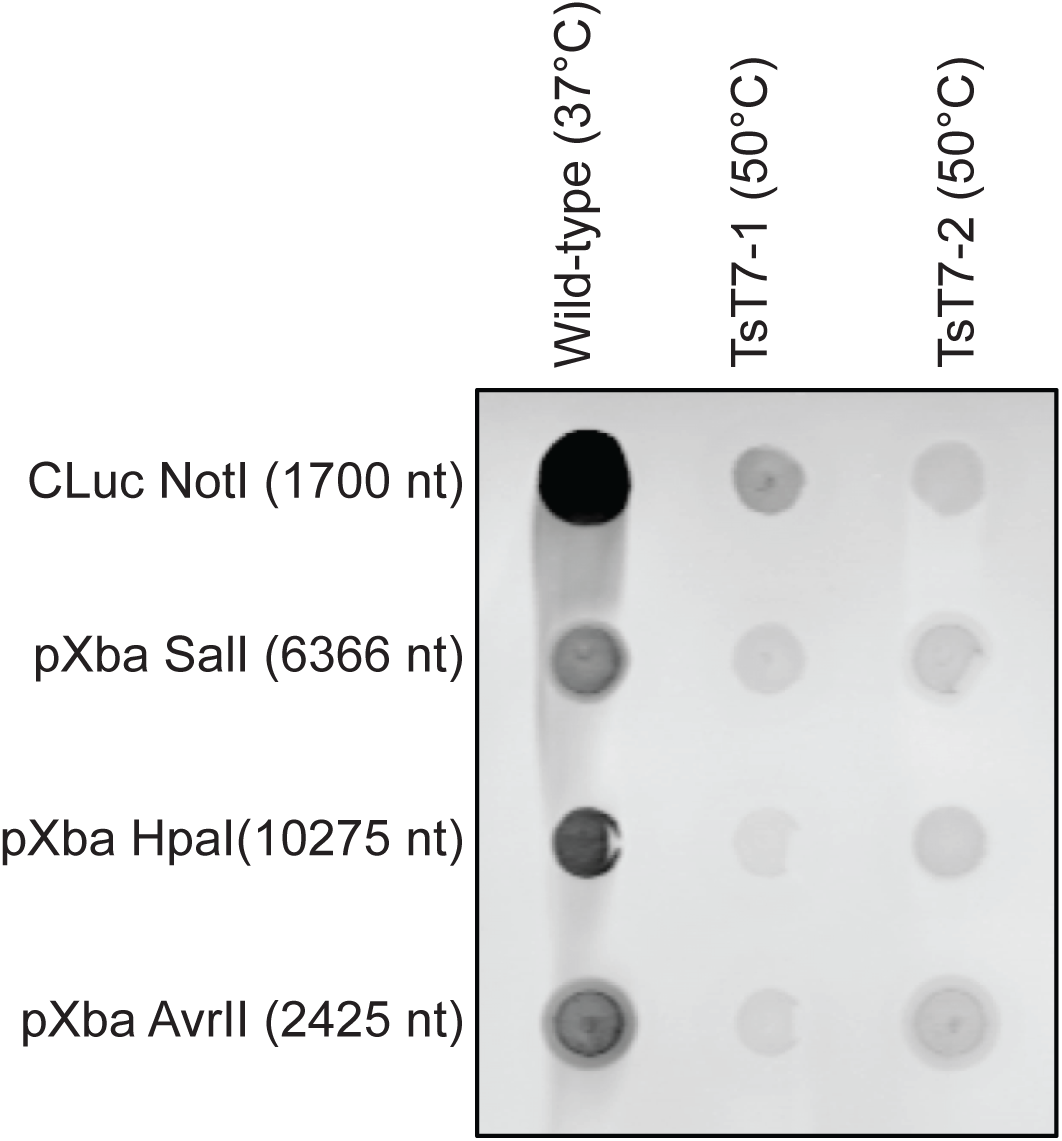
dsRNA immunoblot demonstrating reduction in signal when *in vitro* transcription is performed with TsT7-1 and TsT7-2 at 50°C for multiple templates. Immunoblot on *in vitro* transcription reactions performed on templates with varying length and sequence at the 3’-end with J2 antibody (1:5000; Scicons). *In vitro* transcription reactions were performed with either wild-type T7 RNAP at 37°C, TsT7-1 at 50°C or TsT7-2 at 50°C.

**Supplementary Figure S8.**
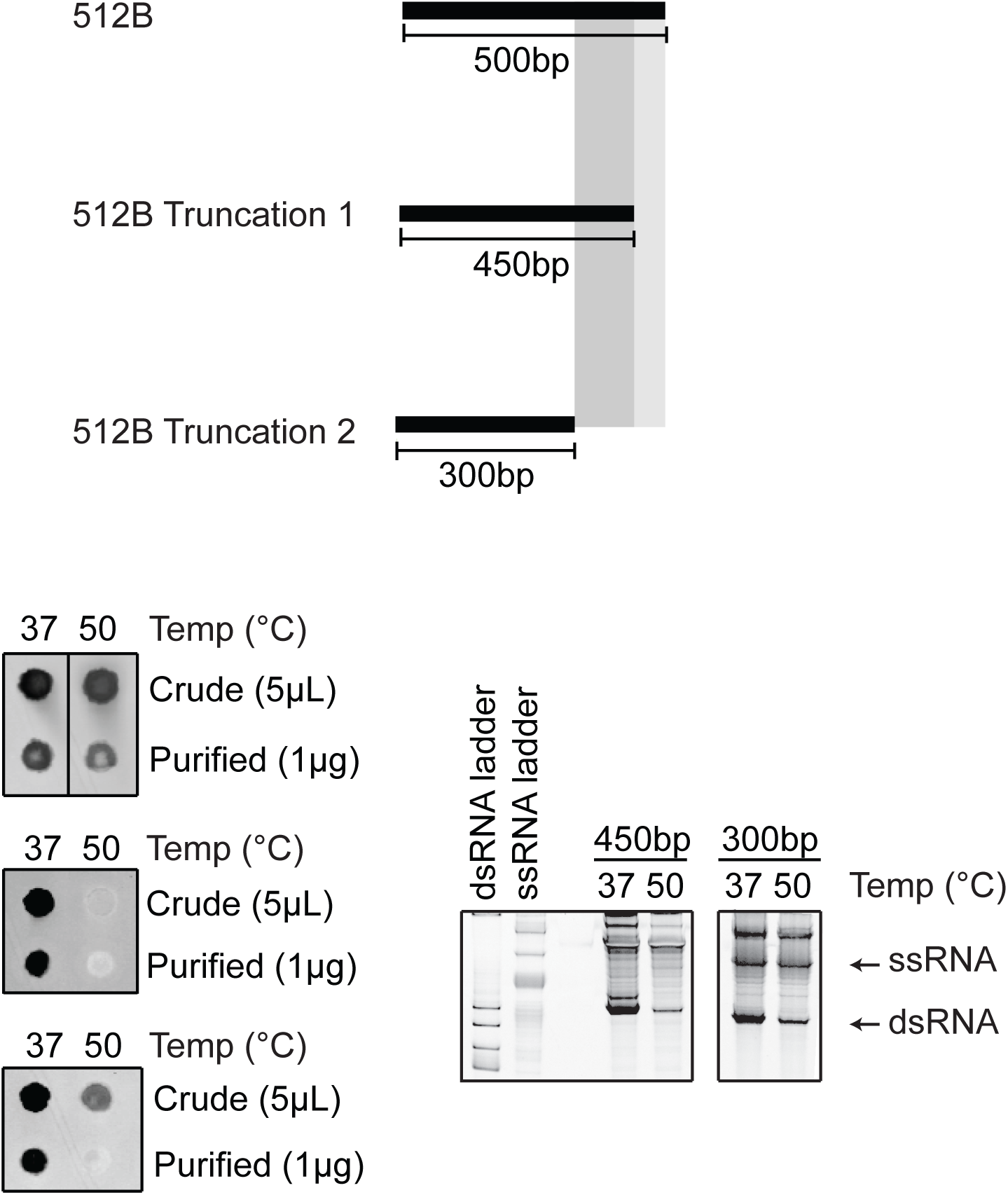
Truncation of the 3’-end of the 512B DNA templates results in reduction of the antisense RNA by-product formation. Immunoblot (with J2 antibody; 1:5000; Scicons) and native gel electrophoresis analyses of *in vitro* transcription reactions performed on 512B template with 3’-end truncations (50 and 200 base pairs). *In vitro* transcription reactions were performed with TsT7-1 at 37°C or 50°C.

**Supplementary Figure S9.**
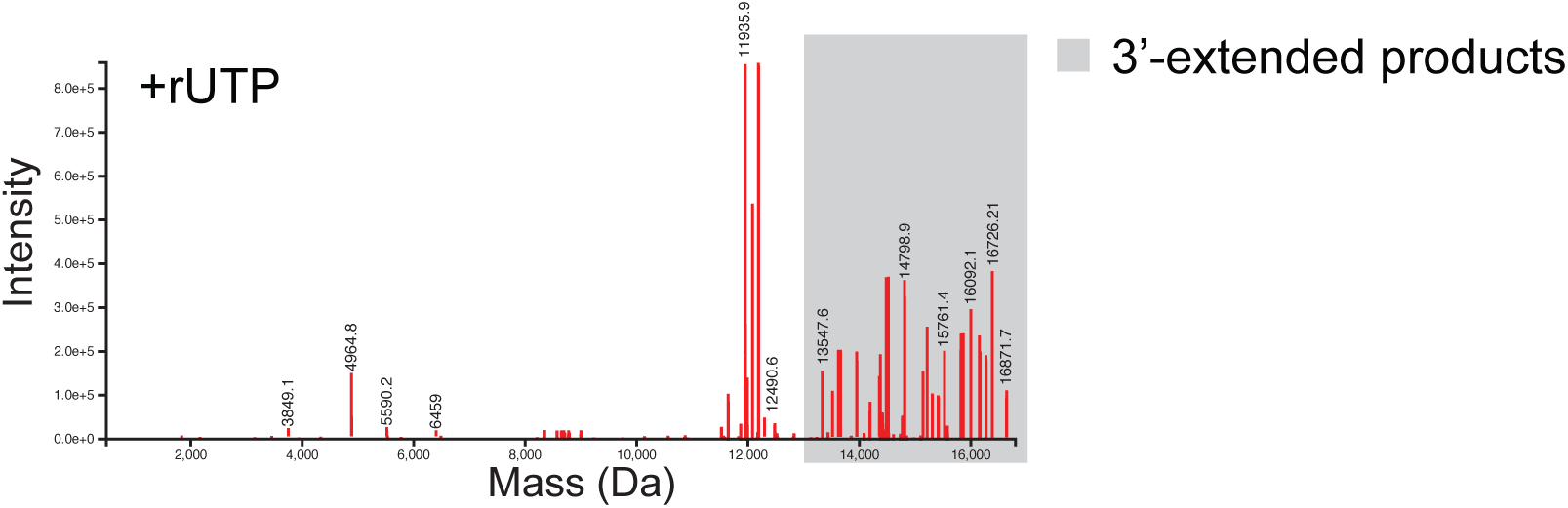
Intact mass spectrometry analysis of IVT Template short-1 with poly-A tail. *In vitro* transcription reactions were performed with wild-type T7 RNAP at 37°C for one hour on IVT Template short-1 templates without or with a poly-A tail (10 base pairs).

